# Essential role of the CCL2-CCR2 axis in Mayaro virus-induced disease

**DOI:** 10.1101/2023.07.21.550077

**Authors:** Franciele Martins Santos, Victor Rodrigues de Costa Melo, Simone de Araújo, Carla Daiane Ferreira de Sousa, Thaiane Pinto Moreira, Matheus Rodrigues Gonçalves, Anna Clara Paiva Menezes dos Santos, Heloísa Athayde Seabra Ferreira, Pedro Augusto Carvalho Costa, Breno Rocha Barrioni, Paula Bargi-Souza, Marivalda de Magalhães Pereira, Maurício Lacerda Nogueira, Danielle da Glória Souza, Pedro Pires Goulart Guimarães, Mauro Martins Teixeira, Celso Martins Queiroz-Junior, Vivian Vasconcelos Costa

## Abstract

Mayaro virus (MAYV) is an emerging arbovirus member of the *Togaviridae* family and *Alphavirus* genus. MAYV infection causes an acute febrile illness accompanied by persistent polyarthralgia and myalgia. Understanding the mechanisms involved in arthritis caused by alphaviruses is necessary to develop specific therapies. In this work, we investigated the role of the CCL2/CCR2 axis in the pathogenesis of MAYV-induced disease. For this, WT C57BL/6J and CCR2^-/-^ mice were infected with MAYV subcutaneously and evaluated for disease development. MAYV infection induced an acute inflammatory disease in WT mice. The immune response profile was characterized by an increase in the production of inflammatory mediators, such as IL-6, TNF and CCL2. Higher levels of CCL2 at the local and systemic levels, was followed by significant recruitment of CCR2^+^ macrophages and a cellular response orchestrated by these cells. CCR2^-/-^ mice showed an increase in CXCL-1 levels, followed by a replacement of the macrophage inflammatory infiltrate by neutrophils. Additionally, absence of the CCR2 receptor protected mice from bone loss induced by MAYV. Accordingly, the silencing of CCL2 chemokine expression *in vivo* and the pharmacological blockade of CCR2 promoted a partial improvement in disease. Cell culture data support the mechanism underlying MAYV’s bone pathology in which: i) MAYV infection promoted a pro-osteoclastogenic microenvironment mediated by IL-6, TNF and CCL2 and ii) migration of osteoclast precursors was dependent on the CCR2/CCL2 axis. Overall, these data contribute to the understanding of the pathophysiology of MAYV infection and to the identification future of specific therapeutic targets in MAYV-induced disease.

**Importance:** This work demonstrates the role of the CCL2/CCR2 axis in MAYV-induced disease. Infection of WT C57BL/6J and CCR2^-/-^ mice was associated with high levels of CCL2, an important chemoattractant involved in the recruitment of macrophages, the main precursor of osteoclasts. In the absence of the CCR2 receptor there is a mitigation of macrophage migration to the target organs of infection and protection of these mice against bone loss induced by MAYV infection. Much evidence has shown that host immune response factors contribute significantly to the tissue damage associated to alfavirus infections. Thus, this work highlights molecular and cellular targets involved in the pathogenesis of arthritis triggered by MAYV, and identifies novel therapeutic possibilities directed to the host inflammatory response unleashed by MAYV.

## Introduction

Mayaro virus (MAYV) is an endemic arthritogenic arbovirus that is widely neglected in South and Central American countries. MAYV belongs to the *Togaviridae* family and the genus *Alphavirus*, which also includes Chikungunya (CHIKV), Ross River (RRV) and Sindbis (SINV) viruses. These viruses are enveloped and have a positive-polarity single-stranded RNA genome (1). MAYV is responsible for sporadic outbreaks of acute febrile illness, particularly in regions around the Amazon basin where it is maintained in wild and rural cycles. *Haemagogus* mosquitoes and nonhuman primates are considered the main vectors and hosts, respectively (2).

Mayaro Fever (MF) is clinically characterized by fever, rash, retro-orbital pain, headache, arthralgia and myalgia. As with CHIKV infection, highly debilitating and persistent arthralgia is a hallmark of MAYV infection, which primarily affects the joints of the wrists, elbows, knees, and small joints of the hands and feet (3–6). Persistent arthralgia occurs in more than 50% of MF cases, which can last from months to years, resulting in large socioeconomic impact (3). The mechanisms associated with MAYV arthralgia pathogenesis are poorly understood. Meanwhile, host inflammatory immune-responses seems to play a key role in the development and progression of alphavirus-induced diseases (7).

Macrophages and their pro-inflammatory products, such as CCL2, TNF and INF-γ are considered the main critical factors in the development of RRV and CHIKV-induced myositis and arthritis (8–10). Additionally, IL-6 and the chemokines CCL2, CCL8 and CCL7 have been reported to be involved in osteoclastogenesis during alphaviral infections (11, 12). Accordingly, a recently proposed animal model for MAYV has shown that MAYV infection induces significant inflammatory responses mediated by TNF, IL-6, INF-γ and CCL2, similar to other alphaviruses (13). Noteworthy, elevated CCL2 levels have been detected in the synovial fluid of patients infected with RRV and CHIKV (14, 15) as well as in the serum of patients with MF during the acute phase of disease (16).

The CCL2-CCR2 axis is responsible for modulating monocyte/macrophage recruitment in several chronic inflammatory diseases including atherosclerosis, cancer, and arthritis (17). In inflammatory sterile conditions such as rheumatoid arthritis (RA) and osteoarthritis (OA), elevated CCL2 levels were detected in blood, synovial fluid and synovial tissue of patients. Furthermore, studies in animal models of osteoarthritis have demonstrated that the CCL2/CCR2 pathway is primarily responsible for the recruitment of monocytes, establishment of pain and associated-tissue damage (18–20). Accordingly, in infectious arthritis models, such as the one induced by CHIKV, CCR2 knockout mice presented more severe arthritis that was associated with an intense neutrophil infiltrate. In contrast, inhibition of CCL2 was associated with a reduction in arthritis, myositis and bone loss in mice infected with CHIKV and RRV (12, 21).

Thus, the aim of the present study was to investigate the role of the CCL2/CCR2 axis in the pathogenesis of MAYV-induced disease. Our results demonstrated that MAYV induced an acute inflammatory disease, characterized by local edema, hypernociception and myositis in both WT and CCR2^-/-^ mice. Meanwhile, viral loads analysis revealed reduction of viable virus titers in most analyzed organs of CCR2^-/-^ mice. Interestingly, in the absence of CCL2/CCR2 signaling, there was protection from bone loss induced by MAYV. *In vitro,* our data support the mechanism underlying MAYV’s bone pathology in which infection promotes a pro-osteoclastogenic microenvironment mediated by IL-6, TNF and CCL2 production as well as migration of osteoclast precursors in a CCL2/CCR2 dependent-manner.

## Results

### Characterization of MAYV-induced disease in C57BL/6J mice

First a characterization of the MAYV-induced disease in young C57BL/6J mice was performed. To that end, 4-week-old mice were infected in the right hind paw with 1×10^6^ PFU of MAYV and signs of disease, including hypernociception, viral loads in several organs and tissue damage during the course of infection were evaluated. MAYV induced significant hind paw edema and reduced the nociceptive threshold as early as with 1-day post-infection (PI) (FIG. 1A and 1B). Hind paw edema peaked at day 5PI and persisted for up to 15 days PI (FIG. 1A). Meanwhile, the nociceptive threshold remained low for up to 28 days upon MAYV inoculation, returning to baseline levels at day 35. These clinical signs were associated with significant inflammatory damage in hind paw tissues (FIG 1C and 1D). Indeed, MAYV induced infiltration of inflammatory cells to the site of the infection, which associated with myositis and loss of tissue architecture specially at day 7 PI (FIG. 1C, 1D v and vi). At days 14 and 21 PI there was still presence of inflammatory infiltrate, but the onset of tissue recovery was already evident, as indicated by the presence of regenerating muscle fibers (FIG. 1D viii and x). Accordingly, MAYV infected, replicated and spread to different tissues, including the contralateral ankle joint and quadriceps femoris muscle, persisting significantly in ankle joints, right quadriceps and paw for up to 7 days PI (FIG 1E).

**Figure 1.**
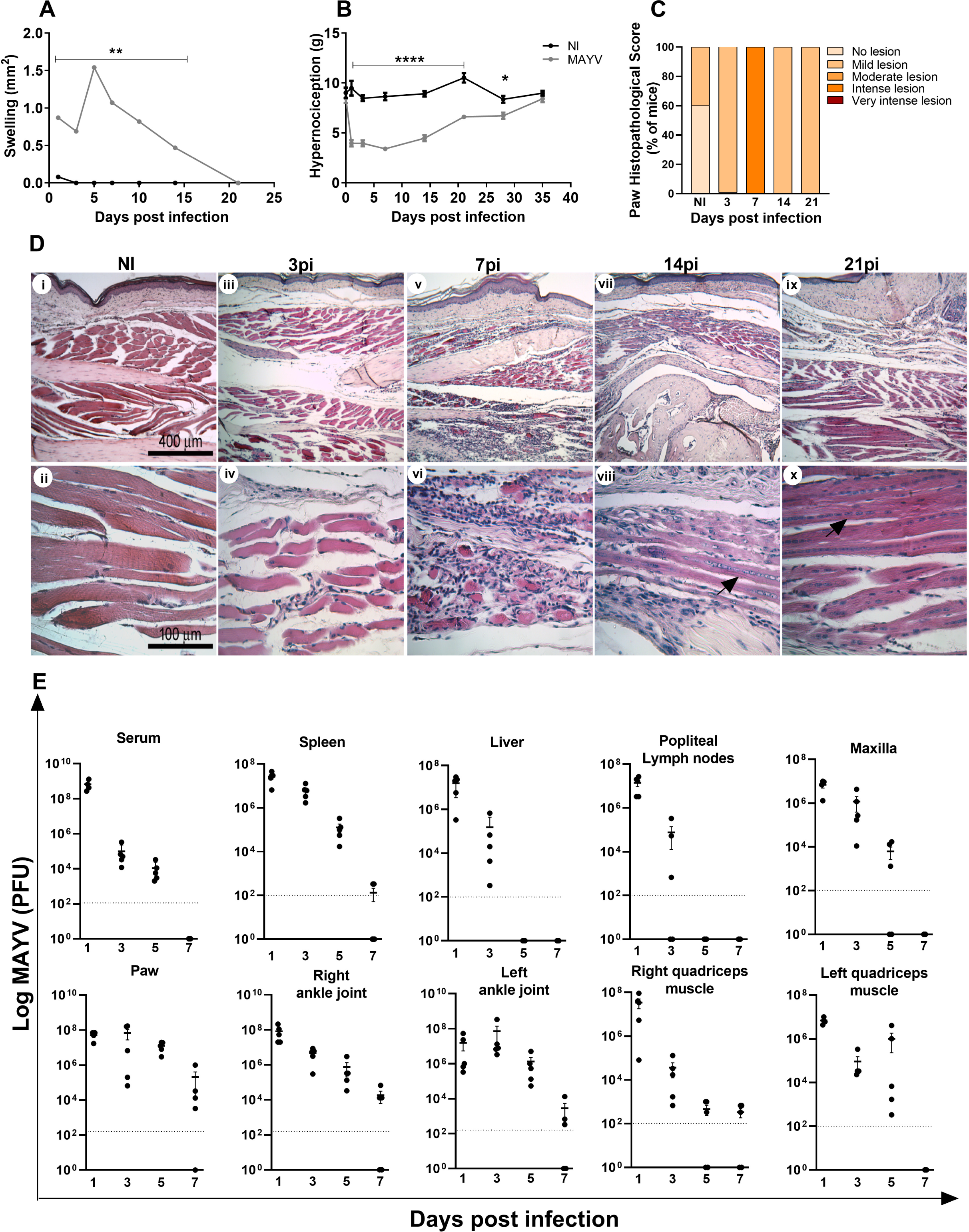
MAYV-induced disease in C57BL/6J mice. 4-week-old mice were infected through the rear right footpad with 30µl containing 10^6^ PFU or were mock infected with PBS 1X. (A) Edema measurements were performed in the infected paw (right paw) with the aid of a caliper. (B) Hypernociception threshold assessment was performed using the adapted von Frey method. Statistics were performed with two-way ANOVA, Turkey’s multiple comparisons test (*P ≤ 0,05; **P < 0.0012; ****P < 0.0001). (C) Histopathological score comprising inflammatory muscle damage. At days 3, 7, 14 and 21 PI mice were euthanized and ankle tissues removed, paraffin embedded and 5 µm sections were generated and stained with HE. Each mouse sample received a score as follows: No lesion (0); Mild lesion (1 and 2); Moderate lesion (3 and 4); Intense lesion (5 and 6) and Very intense lesion (7). The panel D shows negative control (i and ii); MAYV infection at day 3PI (iii and iv); day 7PI (v and vi); day 14PI (vii and viii); day 21PI (ix and x). Dark arrows indicate regenerating muscle fibers. Images i, iii, v, vii and ix (100X magnification, 400 μm bar); images ii, iv, vi, viii and x (400X magnification, 100 μm bar). (E) At days 1, 3, 5 and 7 PI the serum and tissues of interest were harvested, homogenized and the titers of infectious virus was determined by plaque assay on Vero cells. Analyses are representative of at least five mice per group.

MAYV infection induced a marked increase in local (FIG 2A) and systemic (FIG 2E-H) CCL2 levels, which remained high up to 14 days PI in the paw, maxilla, quadriceps muscle and spleen. The CCL5 chemokine was detected in quadriceps and paw tissues (FIG 2B and I). The levels of the pro-inflammatory cytokines IL-6 and TNF increased only in the footpad at day 7PI (FIG 2C and D).

**Figure 2.**
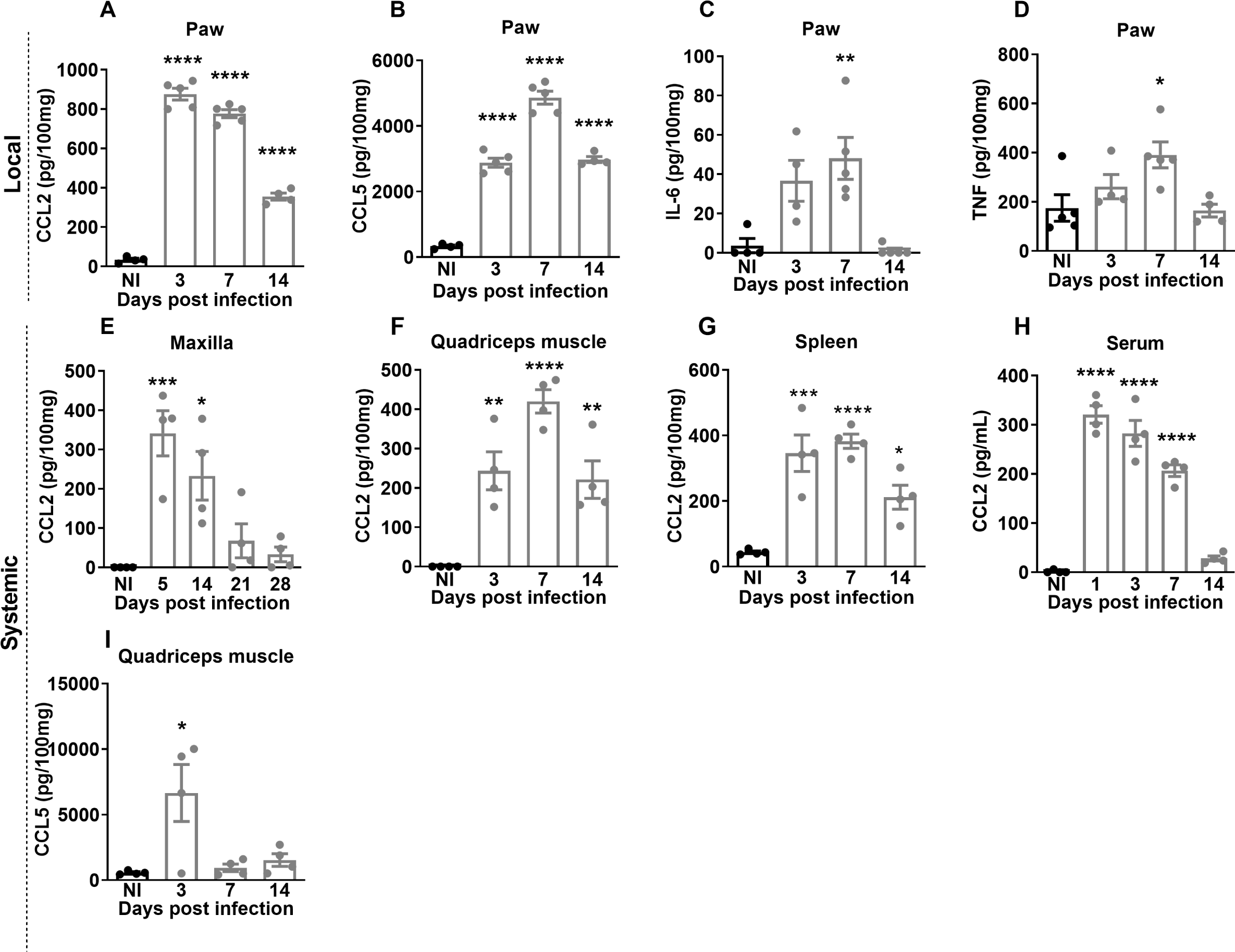
MAYV induces the production of local and systemic inflammatory mediators in C57BL/6J mice. 4-week-old WT were infected through the right hindpaw with 10^6^ PFU of MAYV. NI group was injected with 1X PBS. On days 3, 7 and 14 PI the paws were collected and processed for analysis of the local cytokine profile. The paw samples showed high levels of chemokines CCL2 (A) and CCL5 (B) and inflammatory cytokines IL-6 (C) and TNF (D). For the analysis of the systemic profile of cytokines and chemokines, the quadriceps muscle, serum and spleen were collected on days 3, 7 and 14PI and the maxilla was collected on days 5, 14, 21 and 28PI. The presence of CCL2 chemokines was detected in the maxilla (E), quadriceps muscle (F), spleen (G) and serum (H) and CCL5 in the quadriceps muscle (I). Statistics were performed with one-way ANOVA, Turkey’s multiple comparisons test (*P value ≤0.05; **P value<0.01; ***P value<0,001; ****P value<0.0001). The asterisk indicates the difference between the time points of the infection kinetics and the uninfected group (NI). Analyses are representative of at least four mice per group.

### The CCL2/CCR2 axis plays an important role in the development of MAYV-induced disease

To understand the role of the CCL2/CCR2 axis in the pathogenesis of MAYV, 4-week-old WT and CCR2^-/-^ mice were infected with MAYV to evaluate disease development. Differently from what was observed in WT animals, CCR2^-/-^ mice presented delayed paw edema, peaking at day 10 PI (FIG. 3A). The hypernociception threshold reduced in both WT and CCR2^-/-^ infected mice at day 3 PI (FIG. 3B), with no difference between the groups. However, from the 7^th^ day PI on, CCR2^-/-^ animals recovered and returned to their baseline levels, while WT animals recovered to baseline levels at day 21 PI. As seen in WT animals, MAYV was also detected in the serum and various organs and tissues of the CCR2^-/-^ mice. However, viral titers were significantly lower than in WT in the serum, spleen, liver, right ankle joint and quadriceps muscle, left quadriceps muscle, paw and maxilla (FIG. 3C and D). MAYV-induced inflammation in CCR2^-/-^ mice started at 3 days PI, with the peak on the 7^th^ and 14^th^ days post-virus inoculation (FIG. 4A vii, viii, ix and 4B). Comparing to WT mice (ii-v), the inflammatory process was less intense in CCR2^-/-^ animals (vii-x). On the other hand, it persisted longer in the CCR2^-/-^ mice (FIG. 4A vii-x), as the presence of degenerating fibers and myophagocytosis was still observed at days 14 and 21 post MAYV inoculation (FIG. 4A ix and x). Tissue recovery started at 21 days PI with reduction of inflammatory cells and regeneration of injured tissue, whereas in WT animals this tissue recovery started earlier with 14 days PI (FIG. 4A iv and x).

Our results so far demonstrate that MAYV-infected CCR2**^-/-^** mice present reduced clinical and histological signs of inflammation (FIG 3A-B and 4A-B). Regarding inflammatory mediators in the paw, no significant differences in the expression kinetics of CCL3 (FIG. 4C) was observed when comparing WT *versus* CCR2^-/-^ mice. Meanwhile, CCL5 levels increased at 7 days PI (FIG. 4D), TNF levels reduced at 14 days PI (FIG. 4E), while CCL2 levels were higher at days 7 and 14 PI in knockout than in WT mice (FIG. 4F). Interestingly, the chemokine CXCL-1, a promoter of neutrophil recruitment, was significantly elevated at day 7 PI in knockout mice compared to WT animals (FIG. 4G).

**Figure 3.**
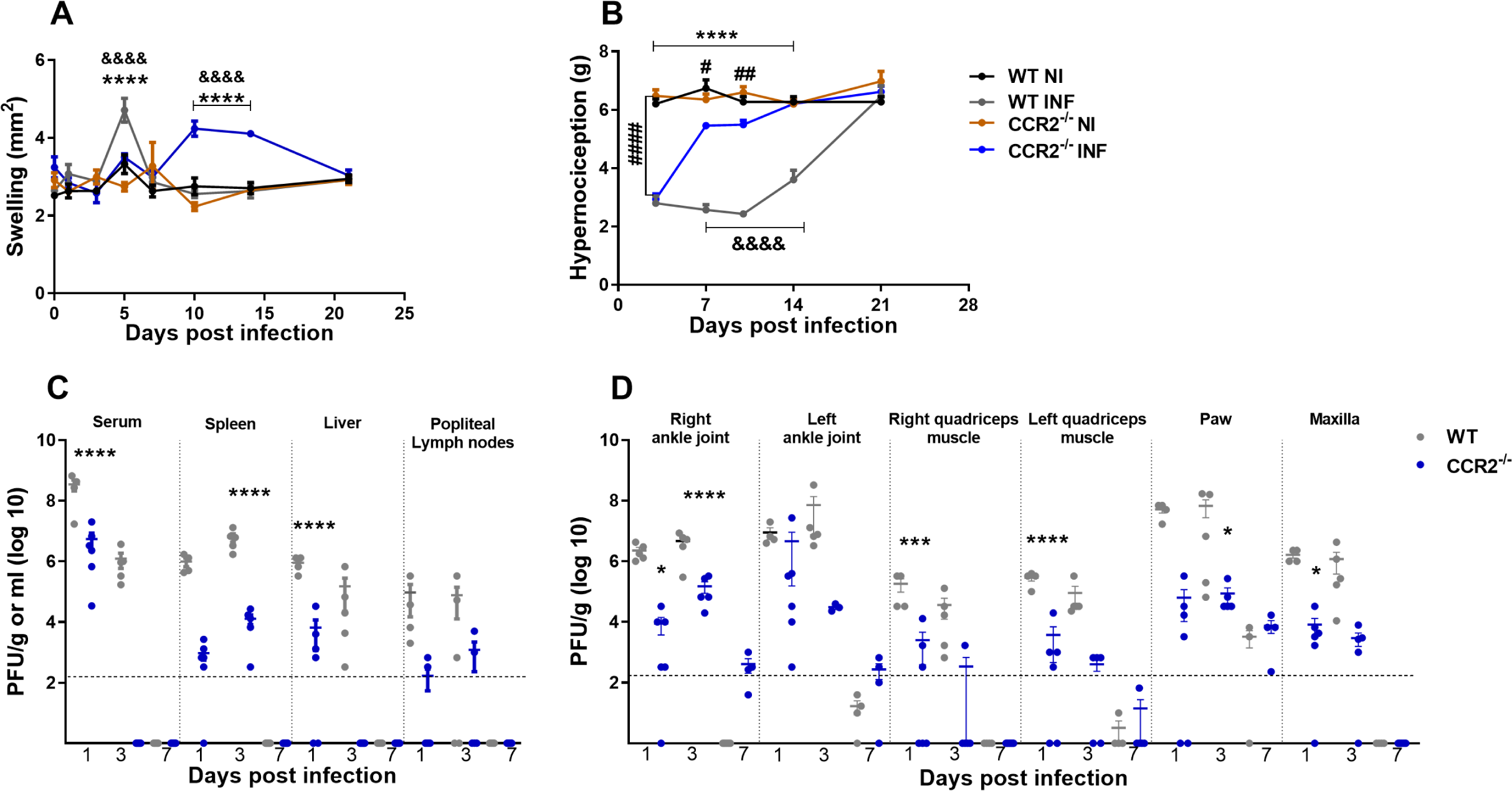
MAYV Infection in CCR2^-/-^ mice. 4-week old WT and CCR2^-/-^ mice (C57BL6 background) were infected via the rear right footpad with 10^6^ PFU of MAYV. Mock group was inoculated with PBS 1X. (A) Plantar edema measurement. Edema was measured with a caliper on the infected paw on 0-, 1-, 3-, 5-, 7-, 10-, 14- and 21-days PI. (B) Measurement of the hypernociception threshold. Statistics were performed with two-way ANOVA, Turkey’s multiple comparisons test (*\*P value ≤0.05; **P value<0.01; ***P value<0,001; ****P value<0.0001*). * difference between WT INF and WT NI; # Difference between CCR2^-/-^ INF and CCR2^-/-^ NI; & difference between WT and CCR2^-/-^. (C and D) At 1-, 3- and 7-days PI serum and tissues of interest were harvested, homogenized and the titers of infectious virus determined by plaque assay on Vero cells. Statistics were performed with student *t*-test at each timepoint (*\*P value ≤0.05; **P value<0.01; ***P value<0,001; ****P value<0.0001*). *Difference between WT and CCR2^-/-^. Analysis are representative of at least five mice per group (Mean ± SEM).

**Figure 4.**
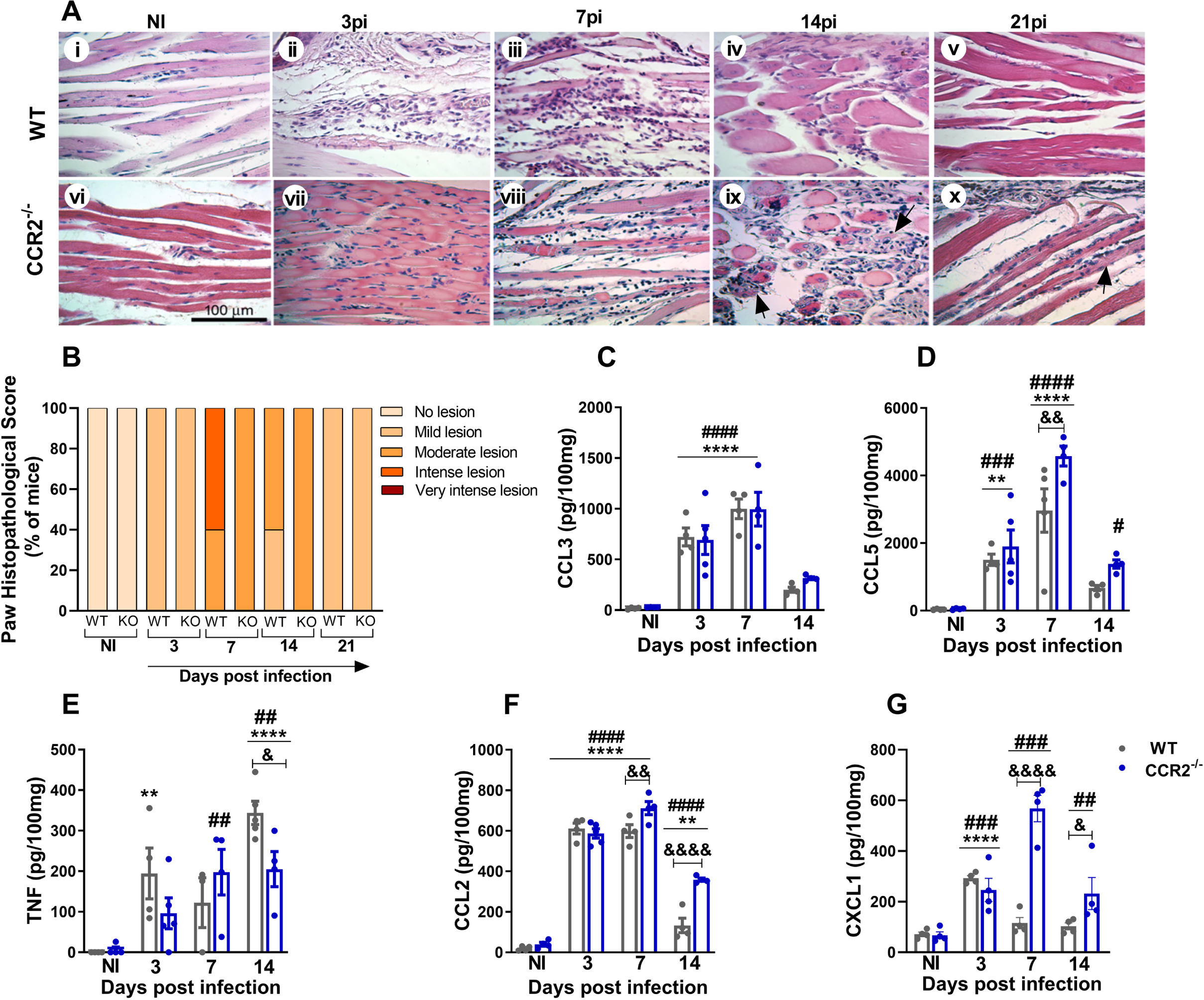
MAYV induces moderate and prolonged inflammation in CCR2^-/-^ mice. (A) At 3-, 7-, 14- and 21-days PI mice were euthanized and ankle tissues removed, paraffin embedded and 5 µm sections were generated and stained with HE. Images i-x (400X magnification, 100 μm bar). (i-v) Panels show WT negative control (i) and MAYV infection at 3 (ii), 7 (iii), 14 (iv) and 21 (v) days PI. Corresponding endpoints are shown for CCR2^-/-^ samples (vi-x). Black arrows indicate suggestive areas of myophagocytosis, as denoted by degenerating muscle fibers surrounded by leukocytes. (B) Paw histopathological score. No lesion (0); Mild lesion (1 and 2); Moderate lesion (3 and 4); Intense lesion (5 and 6) and Very intense lesion (7). (C-G) Kinetics of expression of the inflammatory mediators CCL3 (C), CCL5 (D), TNF (E), CCL2 (F) and CXCL1 (G) in the mice’s paws on days 3, 7 and 14 PI. Statistics were performed with two-way ANOVA, followed by Tukey’s multiple comparisons test (*\*P value ≤0.05; **P value<0.01; ***P value<0,001; ****P value<0.0001*) * difference between WT INF and WT NI; # Difference between CCR2^-/-^ INF and CCR2^-/-^ NI; & difference between WT and CCR2^-/-^. Analysis are representative of at least five mice per group (Mean ± SEM).

### MAYV induces bone loss in a CCR2-dependent manner

The inflammatory response induced by MAYV infection was characterized by a significant pro-osteoclastogenic milieu, as demonstrated by increased local expression of IL-6, TNF and CCL-2, and systemic overexpression of CCL2. To assess the impact of MAYV infection on bone tissue, samples of tibia were submitted to microtomography (microCT - μCT) analysis at 21-days PI. Bone parameter data comparing WT and CCR2^-/-^ mice were expressed as delta (infected minus non-infected for each sample) to eliminate the bias of the baseline difference in bone mass that exists between these animals. MAYV induced significant bone loss in the distal tibial metaphysis region of WT mice, in contrast to CCR2^-/-^ mice (FIG. 5). There was a significant reduction in bone volume (FIG. 5A) in the tibia of WT infected mice, as well as in the trabecular number (FIG. 5B), accompanied by increase in the porosity (FIG. 5C and F i-ii). In contrast, bone homeostasis in CCR2^-/-^ animals was not affected by MAYV infection. There was no significant bone resorptive phenotype in the tibia of CCR2^-/-^ mice (FIG. 5A, B, C, D and F).

**Figure 5.**
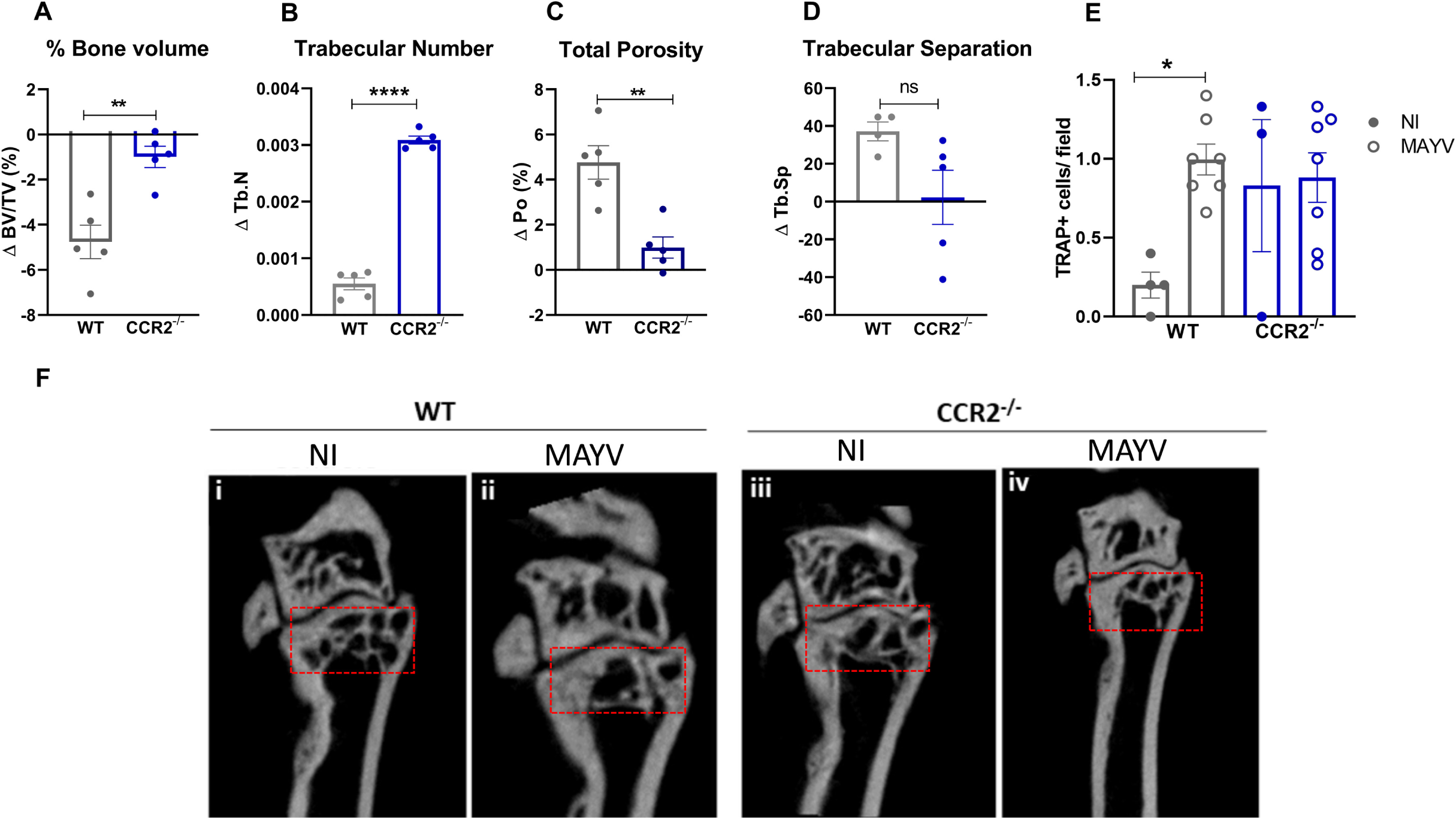
CCR2 ^-/-^ mice are resistant to bone resorption induced by MAYV. 4-week old WT and CCR2^-/-^ mice (C57BL6 background) were infected via the rear right footpad with 10^6^ PFU of MAYV. Mock group was inoculated with PBS 1X. At day 21 PI animals were euthanized, and the tibia removed for microCT analysis. (A) Trabecular bone volume fraction (BV/TV (%)), (B) Trabecular Number (Tb.N), (C) Porosity in the tibia (Po (%)), (D) Trabecular Separation (Tb.Sp). Statistics were performed with Student’s unpaired t test (**P value <0.01). Analysis are representative of at least five mice per group. (E) Number of Trap^+^ cells detected in the distal tibial metaphysis region per microscopic 400x field. Statistics were performed with one-way ANOVA, followed by Tukey’s multiple comparisons test (*P value <0,05). (F) 2D reconstruction of the distal tibia samples. Red box: region of analysis.

To confirm the increase in bone resorption activity in the tibia, we performed TRAP (tartrate-resistant acid phosphatase) staining, an enzyme expressed by osteoclasts during bone resorption. The number of TRAP+ cells was significantly increased in WT-infected animals, while CCR2^-/-^ mice, despite increased number of TRAP+ cells in the basal condition, did not present higher numbers after infection (FIG. 5E). These data indicate that the CCR2 receptor contributes to the bone resorptive phenotype observed during MAYV infection.

### Absence of CCR2 is associated with a shift of mononuclear infiltrate to polymorphonuclear cells resulting in better disease outcome upon MAYV infection

To determine the pattern of the cellular immune response against MAYV infection, WT and CCR2^-/-^ immune cells were isolated from the hind paw and the cellular profile evaluated by flow cytometry. To create the boolean gate, debris were excluded through the combinations of fluorochromes, remotion of doublets a forward scatter area (FSC-A) versus forward scatter height (FSC-H) gate was used, and then cells were gated in function of time versus FSC-A to avoid a possible interference of flux interruptions. MAYV infection increased the frequency in myeloid cells in both infected groups on day 7 PI, with a greater increase in the WT group (FIG 6A and B). At day 1 PI, WT mice showed a reduction in the populations of dendritic cells (CD11c^+^MHCII^+^CD11b^+^) and resident macrophages (Ly6C^low^F4/80^+^CD11b^+^) (FIG. 6 C and D). WT mice showed a response mainly characterized by monocytes/macrophages, which are suggested to be the main cell population involved in alphaviral pathogenesis (13). An increase in the number of inflammatory monocytes (Ly6C^high^F4/80^+^CD11b^+^) was observed 7 days after MAYV inoculation (FIG. 6E) and approximately 30% of these cells were CCR2^+^ monocytes (FIG 6J). Finally, a 2-fold increase in the population of dendritic cells and resident macrophages (Ly6C^low^F4/80^+^CD11b^+^) was observed at day 7 PI (FIG. 6C and D), and approximately 50% of these macrophages are CCR2^+^ (Ly6C^low^CCR2^+^) (FIG. 6K). Interestingly, no significant increase in neutrophils (Ly6G^+^F4/80^-^CD11b^+^) was observed in WT animals after inoculation of MAYV (FIG. 6F). Finally, activation state of these myeloid cell populations was confirmed by cell surface molecule CD80, on monocytes and dendritic cells (FIG. 6G-I).

**Figure 6.**
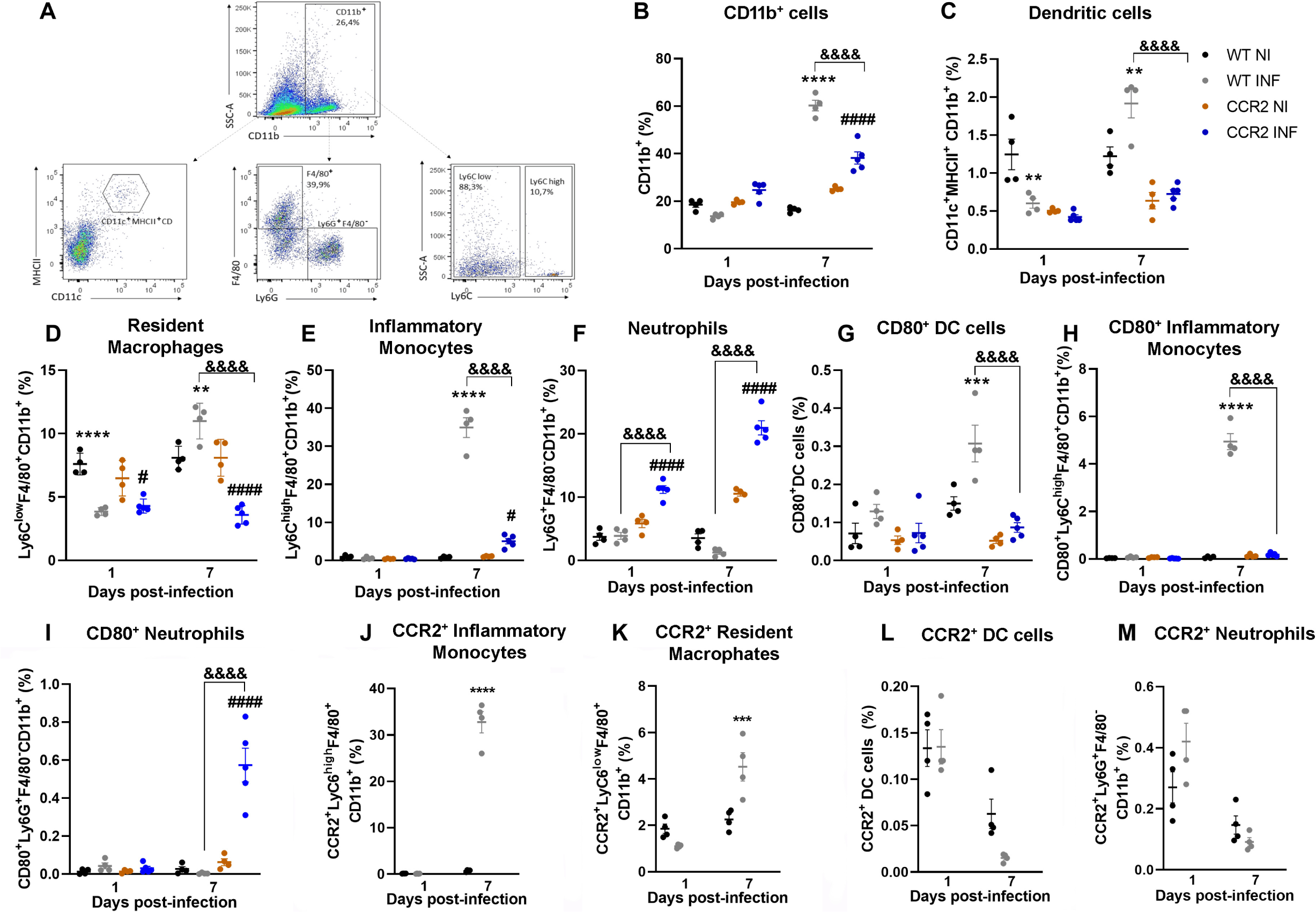
Profile of myeloid immune cells in the paw of MAYV-infected WT and CCR2^-/-^ mice. Flow cytometry analysis of immune cells isolated from the hindpaw of animals infected with MAYV at days 1 and 7 PI. (A) Panel shows gating strategy to define the population of myeloid cells from the single cells. (B) Increase of CD11b^+-^ cells at 7 days PI in WT and CCR2^-/-^ animals. (C) Percentage of dendritic cells (CD11c^+^MHCII^+^CD11b^+^). (D) Percentage of inflammatory monocytes (Ly6C^high^F4/80^+^CD11b^+^). (E) Percentage of resident macrophages (Ly6C^low^F4/80^+^CD11b^+^). (F) Percentage of neutrophils (Ly6G^+^F4/80^-^CD11b^+^). (G) Percentage of dendritic cells expressing CD80. (H) Percentage of inflammatory monocytes expressing CD80. (I) Percentage of neutrophils expressing CD80. Percentage Inflammatory monocytes (J), macrophages (K), dendritic cells (L) neutrophils (M) expressing CCR2. Statistics were performed with two-way ANOVA, followed by Tukey’s multiple comparisons test (*\*P value ≤0.05; **P value<0.01; ***P value<0,001; ****P value<0.0001*). *difference between WT NI and WT INF; # Difference between CCR2^-/-^ NI and CCR2^-/-^ INF; & difference between WT and CCR2^-/-^. Analysis are representative of at least four mice per group (Mean ± SEM).

On the other hand, cellular responses of CCR2^-/-^ mice were characterized by a reduced population of infiltrating inflammatory monocytes/macrophages (FIG. 6E), which was replaced by a significant increase in neutrophils, observed on days 1 and 7 after MAYV inoculation (FIG. 6F). Furthermore, the activation status of these neutrophil population was confirmed by an increase in CD80 positive cells at day 7 PI (FIG. 6I). Finally, both infected-groups revealed an increased percentage of CD4^+^ and CD8^+^ lymphocytes at 7 days PI (FIG. 7A, B and C), while CCR2^-/-^ mice showed a higher percentage of CD4^+^ T cells in comparison to WT. Consequently, activation marker analysis showed an increase in the percentage of CD4+ and CD8+ T cells positive for CD69 and CD44 markers in both infected groups, while CCR2^-/-^ mice showed higher percentage of these cells than WT mice (FIG. 7E-F and H-I). Interestingly, an increase in percentage of T cells CD44^+^/CD62L^+^, a marker for memory cells, was observed exclusively in WT (FIG. 7G and J) mice. Indeed, T lymphocytes from WT mice positive for the CCR2 receptor were observed (FIG. 7K-L).

**Figure 7.**
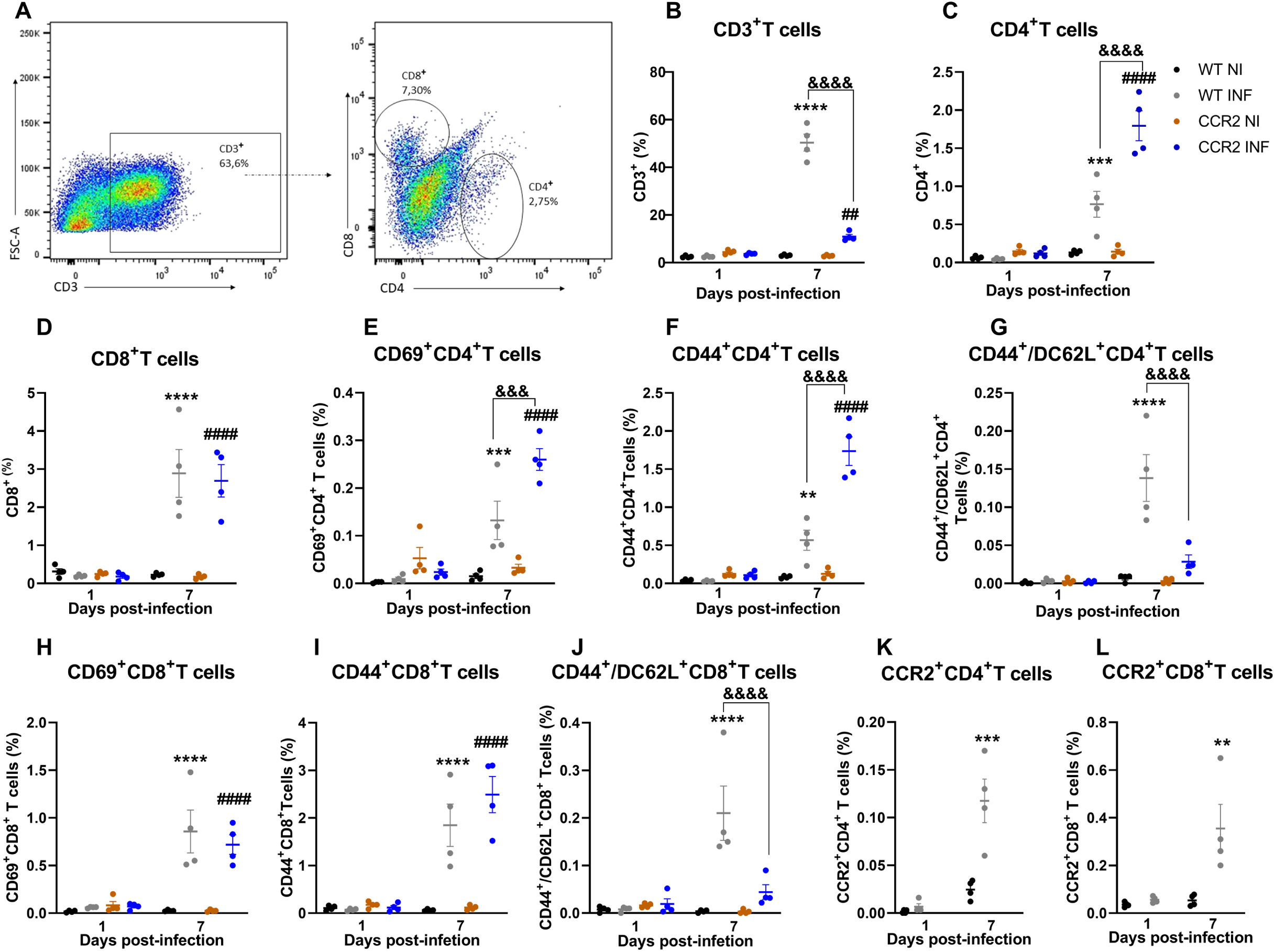
Profile of lymphoid immune cells in the paw of MAYV-infected WT and CCR2^-/-^ mice. Flow cytometry analysis of immune cells isolated from the hindpaw of animals infected with MAYV at days 1 and 7 PI. (A) Panel shows gating strategy to define the population of lymphoid cells from the single cells. % of total cells were used to represent T cell subsets. (B) CD3^+^ cells population. (C) CD4^+^T cells population. (D) CD8^+^ T cells population. (E) Percentage of CD4^+^T cells expressing CD69. (F) Percentage of CD4^+^T cells expressing CD44. (G) Percentage of CD4^+^T cells expressing CD44/CD62L. (H) Percentage of CD8^+^T cells expressing CD69. (I) Percentage of CD8^+^T cells expressing CD44. (J) Percentage of CD4^+^T cells expressing CD44/CD62L. Percentage of CD4^+^T (K) and CD8^+^T (L) cells expressing CCR^-/-^. Statistics were performed with two-way ANOVA, followed by Tukey’s multiple comparisons test (*P value ≤0.05; **P value<0.01; ***P value<0,001; ****P value<0.0001). *difference between WT NI and WT INF; # Difference between CCR2^-/-^ NI and CCR2^-/-^ INF; & difference between WT and CCR2^-/-^. Analysis are representative of at least four mice per group (Mean ± SEM).

Overall, these data show that during MAYV infection, the CCR2 receptor directs the influx and activation of monocytes to the site of infection, which negatively affects the course and severity of the disease. Meanwhile, in the absence of the CCR2 receptor, a differential cellular pattern of cellular infiltration occurs characterized by the presence of neutrophils. This replacement of monocytes by neutrophils in CCR2-/-mice may largely explain the improvement observed in important parameters of the disease, such as hypernociception, bone loss and viral load. Furthermore, the absence of monocytes at the site of infection may be associated with a delay in the onset of edema and recovery from tissue damage.

### Interference in the CCL2-CCR2 axis with two therapeutic approaches showed positive effects on the pathogenesis of MAYV

In complementary experiments, we performed CCL2 gene silencing and pharmacological antagonism of the CCR2 receptor *in vivo* to understand the full role of the CCR2-CCL2 axis in the pathogenesis of MAYV and to explore the therapeutic potential of these molecules. For the silencing, siRNA-CCL2 complexed of polymeric-lipid nanoparticle (NP-siRNA-CCL2) was used to silencing the *ccl2* chemokine expression to support our findings in CCL2/CCR2 pathway (FIG. 8A). We employed the Dynamic Light Scattering (DLS) technique to determine the precise particle size, zeta potential and Polydispersity Index (PDI). DLS measurements showed that the mean diameter of NP-siRNA-CCL2 was 121.3 nm (SD 0.55 nm) and PDI 0.15 (SD 0.01) and zeta potential around -0.51 mV (SD 0.18 mV). Silencing *Ccl2* chemokine expression in WT mice with SiRNA did not induce any effect in paw edema, although it improved the hypernociception thresholds significantly (FIG. 8B and C). Additionally, treatment of WT mice with SiCCL2 also decreased CCL2 levels and *Ccl2*, *Ccl*5, *Ccl7* and *Tnf* expression in hindpaw tissue (FIG. 8D-I). Meanwhile, no difference in the inflammatory infiltrate and histological damage unleashed by MAYV inoculation was observed (FIG. 8 J-K).

**Figure 8.**
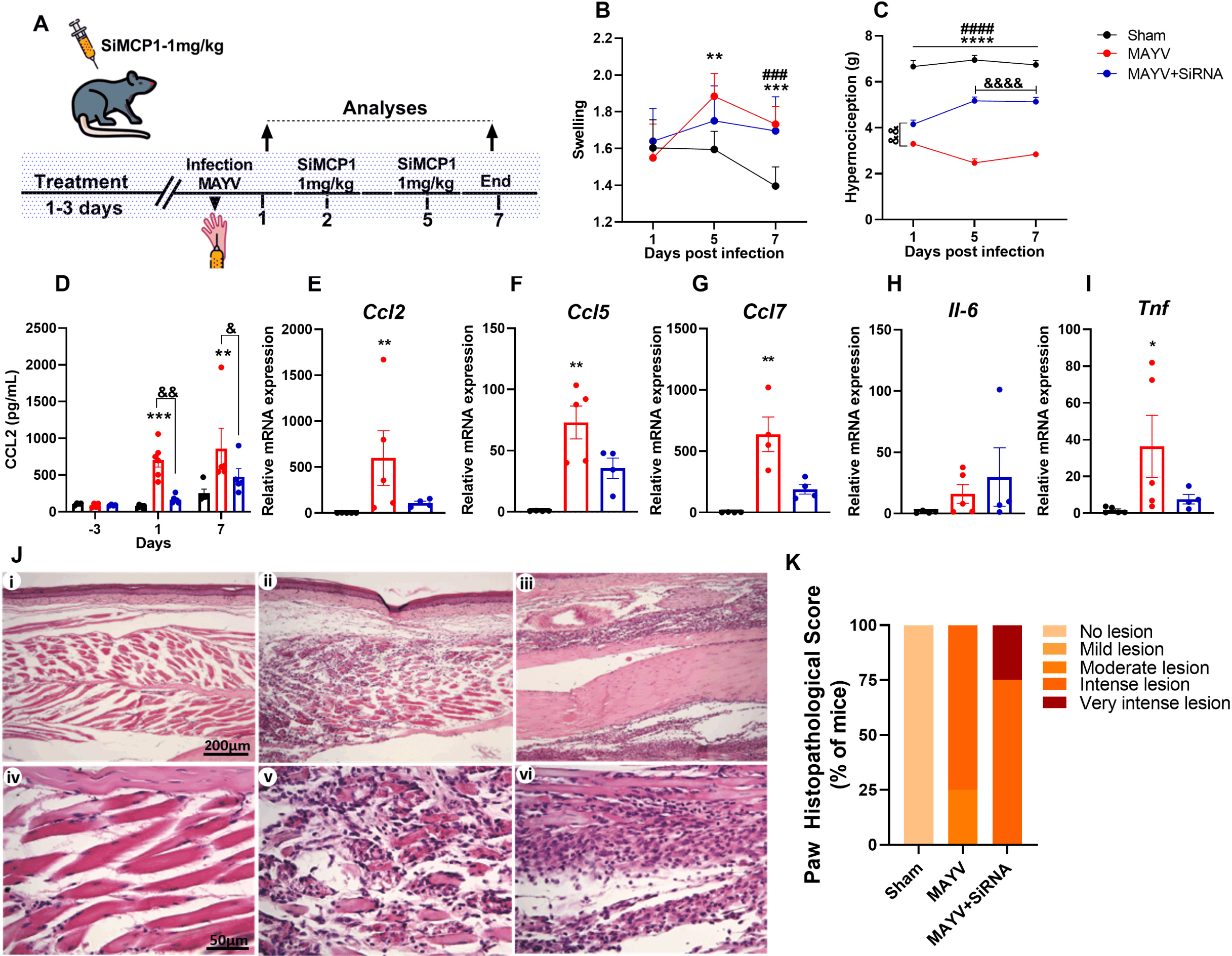
CCL2 chemokine silencing in C57BL6 mice with SiRNA. 4-week old mice were treated for 3 days with NP-siRNA-CCL2 (NP-siRNA-MCP1) before infection and on days 2 and 5 post-infection. Animals were infected via the rear right footpad with 10^6^ PFU or were mock infected with PBS1X. (A) Experimental design. (B) Edema measurements were performed in the infected paw (right paw) with the aid of a caliper on days 2 and 5 post-infection. (C) Hypernociception threshold assessment was performed using the von Frey method. Statistics were performed with two-way ANOVA, followed by Tukey’s multiple comparisons test (*P value ≤0.05; **P value<0.01; ***P value<0,001; ****P value<0.0001). *difference between WT NI and WT INF; # Difference between CCR2^-/-^ NI and CCR2^-/-^ INF; & difference between WT and CCR2^-/-^. (D) Serum CCL2 levels on days -3 (before to infection) and on days 1 and 7 post-infection. (E-I) qRT-PCR analysis of the relative gene expression of the chemokines *Ccl2* (E), *Ccl5* (F), *Ccl7* (G) and cytokines *Il-6* (H) and *Tnf* (I) in the paw. Statistics were performed with Kruskal-Wallis test (*P value ≤0.05; **P value<0.01; ***P value<0,001; ****P value<0.0001). (J) Representative histological micrographies. At day 7 PI mice were euthanized and ankle tissues were removed, paraffin embedded and 5 µm sections were generated and stained with HE. The panel shows negative control (i and iv), animals infected with MAYV (ii and v) animals infected and treated with SiMCP1 (iii and vi). (P) Histopathological paw score.

Additionally, we performed the pharmacological blockade of CCR2 with a CCR2 small antagonist molecule, RS504393. The continuous treatment with RS504393 had no effect on viral load in most of the analyzed tissues, showing a discrete reduction only in the paw (FIG. 9A and B). Also, the CCR2 antagonist did not interfere with the levels of CXCL1 or CCL2, although it reduced the CCL5 concentrations in the paw (FIG. 9C-E). The histopathological analysis revealed a reduction in percentage of mice presenting intense tissue damage and leukocyte infiltration after the treatment (FIG. 9F i-iv and G). In view of the potential of MAYV to induce bone loss and the signaling role of CCR2 during this pathological process, we also investigated whether treatment with RS504393 would protect mice against bone resorption. The µCT results showed that the pharmacological treatment induced a milder protective phenotype than that observed in CCR2^-/-^ mice. The mean values for total bone volume, trabecular separation, porosity and trabecular number had a discrete reduction in mice treated with the CCR2 antagonist, but there was no statistical difference when compared with the vehicle-treated mice (FIG. 9H-K).

**Figure 9.**
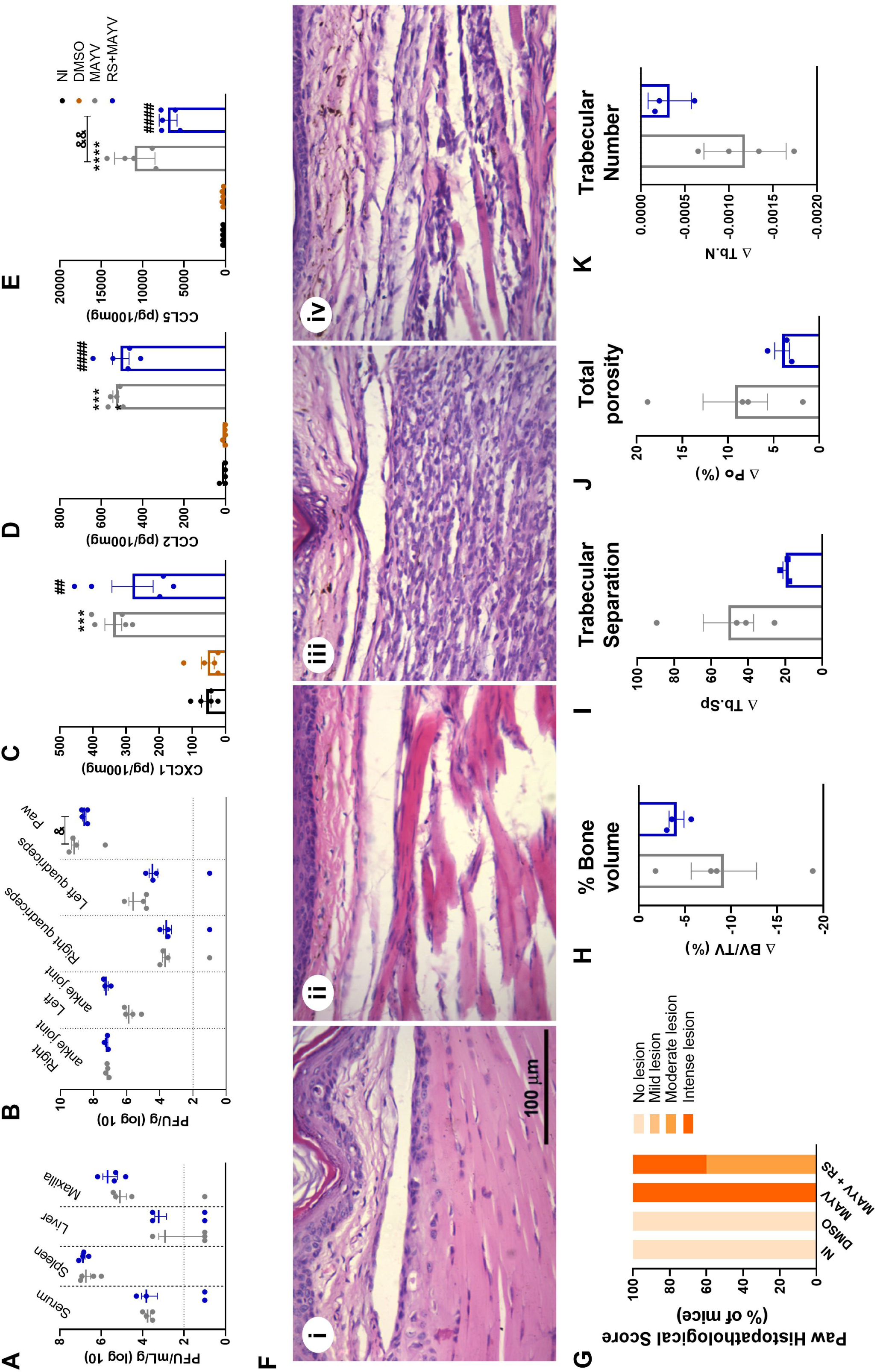
Effect of CCR2 antagonism on MAYV-induced disease. 4-week old mice were treated with RS504393 1 day before infection and every 24 days after infection until the point of analysis. To analyze the effect of RS504393 on viral load the animals were evaluated up to day 2 PI and tissues were collected on day 3 PI (A and B). To evaluate the chemokine profile and the inflammatory process in the paw, the animals were treated until day 6 PI and the paw tissues of interest were collected on day 7 PI. (C) CXCL1, (D) CCL3 and (E) CCL5 levels were evaluated by ELISA. Statistics were performed with on-way ANOVA, followed by Tukey’s multiple comparisons test (*P value ≤0.05; **P value<0.01; ***P value<0,001; ****P value<0.0001). (F) Representative images of paw tissue samples. At day 7 p.i. mice were euthanized and paw tissues were removed, paraffin embedded and 5 µm sections were generated and stained with HE. The panel (i) shows negative control, group DMSO (ii), group infected with MAYV (iii) and group infected and treated with RS504393 (iv). (G) Histopathological paw score. With 20 days of treatment and 21 days PI the mice were euthanized and the tibia was collected for analysis by µCT. (H) % Bone volume (BV/TV (%)), (I) Trabecular Separation (Tb.Sp), (J) Total porosity (Po (%)) and (K) Trabecular Number (Tb.N). Statistics were performed with Student’s Unpaired t test (no significant difference has been detected).

### Response of macrophages, osteoclasts and osteoblasts against MAYV

Bone homeostasis is the result of a balance between osteocyte signaling, osteoblastic bone formation, and osteoclastic bone resorption (34). To understand the mechanism by which MAYV caused bone loss *in vivo*, we investigated the susceptibility and response of macrophages, osteoclasts and osteoblasts after MAYV infection *in vitro*. Infection of primary culture WT and CCR2^-/-^ macrophages with MAYV at MOI of 1 showed that although there was no productive infection (FIG 10A) or loss of viability (FIG 10D), MAYV modulated the production of cytokines by these cells. Macrophage response to infection resulted in IL-6 production only by WT cells (FIG 10B), while TNF was significantly detected in CCR2^-/-^ cells at 72h PI (FIG 10D). Similarly, MAYV was unable to promote a productive infection in primary osteoclasts (RANKL-differentiated) (FIG 10E), but momentarily reduced the viability of the CCR2^-/-^ cells after 24 and 48h of infection (FIG 10H). MAYV infection also induced the production of IL-6 (FIG 10F) and TNF (FIG 10G), two important pro osteoclastogenic mediators, in both WT and CCR2^-/-^ osteoclast cultures. Nevertheless, such IL-6 and TNF production was significantly lower in CCR2^-/-^ osteoclasts when compared to WT cells (FIG 10F, G). To investigate the osteoblast response to infection, bone marrow-derived mesenchymal stem cells (BMSCs) were isolated and followed differentiation protocol for 14 days. BMSCs isolated from WT and from CCR2^-/-^ animals were cultivated to compare the response of these cells *in vitro*. There was no productive MAYV infection in WT and CCR2^-/-^ cells (FIG 10I). Furthermore, no loss of viability (FIG 10K) and changes in the alkaline phosphatase activity in osteoblasts with active CCR2 (NI: 0.224±0.063 *versus* INF: 0.267±0.091, *p* = 0.9999) or CCR2^-/-^ (NI: 0.219±0.046 *versus* INF: 0.218 ±, 0,057, *p* > 0.9999) have been observed. Interestingly, IL-6 was produced exclusively by WT osteoblasts in response to MAYV infection (FIG. 10J), which suggests that the cytokine IL-6 plays, along with TNF, an important role in the mechanism of bone loss induced by MAYV. Similar results were detected when CCR2 was antagonized by RS504393, in which osteoblasts with active CCR2 showed a significant production of IL-6 after infection (NI: 879.73±198.27 pg/mL *versus* INF: 2,966.72±435.27 pg/mL, *p* < 0.0001) when compared to cells with blocked CCR2 (NI: 1,132.73±643.82 pg/mL; INF: 1,250.96±628.95 pg/mL, *p* = 0.9921).

**Figure 10.**
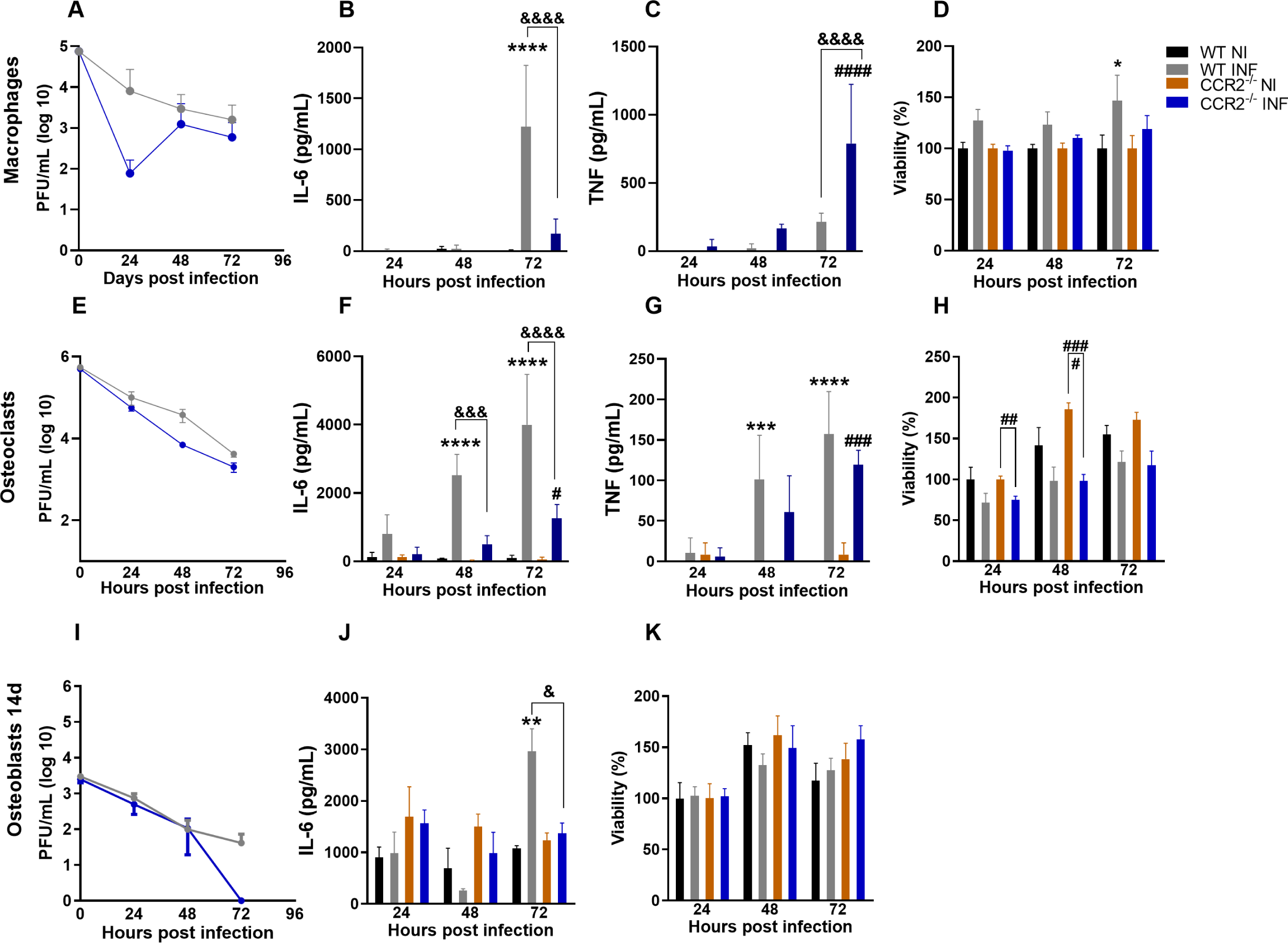
*In vitro* infection of macrophages, osteoclast and osteoblasts by MAYV. (A) MAYV titration in primary macrophages. (B) Quantification of IL-6 and (C) TNF levels in the macrophage culture supernatant. (D) Cell viability of primary culture of WT and CCR2^-/-^ macrophages. (E) MAYV titration in primary osteoclasts. (F) Quantification of IL-6 and (G) TNF levels in the osteoclast culture supernatant. (H) Cell viability of primary culture of WT and CCR2^-/-^ osteoclasts. (I) MAYV titration in the primary culture of osteoblasts from WT and CCR2^-/-^ cells differentiated for 14 days. (J) Quantification of IL-6 in the osteoblast culture. (K) Cell viability of primary osteoblast culture of WT and CCR2^-/-^ cells. Statistics were performed with two-way ANOVA, followed by Tukey’s multiple comparisons test (*P value ≤0.05; **P value<0.01; ***P value<0,001; ****P value<0.0001). *difference between WT NI and WT INF; # Difference between CCR2-/-NI and CCR2-/-INF; & difference between WT and CCR2-/-. Analyses are representative of at least three biological replicates (Mean ± SEM).

In sumary, these data show that the inflammatory response to MAYV infection differs in the presence or absence of CCR2, which is associated with a lower production of inflammatory and pro-osteoclastogenic cytokines, including IL-6 and TNF.

## Discussion

MAYV is an emerging alphavirus in the American continent, that has great potential for urbanization and has recently become an important public health problem in Brazil (22). MAYV, like RRV and CHIKV, is responsible for causing a highly debilitating musculoskeletal inflammatory disease, which includes polyarthralgia/polyarthritis and myalgia. The disease induced by arthritogenic alphaviruses is classified as an immunopathology (7, 23). Immunopathology is being increasingly recognized in many human viral diseases, such as dengue, zika, influenza and more recently in COVID19 (24–26). Therefore, the investigation of the immune response mechanisms that influence the pathogenesis of MAYV infection would allow to design better treatment strategies to limit inflammation and tissue damage driven by inflammation (7). The major findings of the present study can be summarized as follows: i. Infection of C57BL/6J mice induced an acute inflammatory disease, characterized by local edema, hypernociception, myositis, replication in target organs, production of several inflammatory mediators and bone resorption; ii) Absence of CCR2 receptor has a protective effect on MAYV infection, showed by signs of milder and delayed disease manifestation, reduced viral loads in several organs, less tissue damage and absence of bone loss; iii) The inflammatory profile of cell infiltrate of WT mice is predominantly composed by macrophages, whereas in CCR2^-/-^ animals it has been replaced by neutrophils; iv) MAYV has the potential to modulate the intrinsic ability of osteoblasts, osteoclasts and macrophages to produce pro-inflammatory cytokines related with bone resorption (FIG. 11).

**Figure 11.**
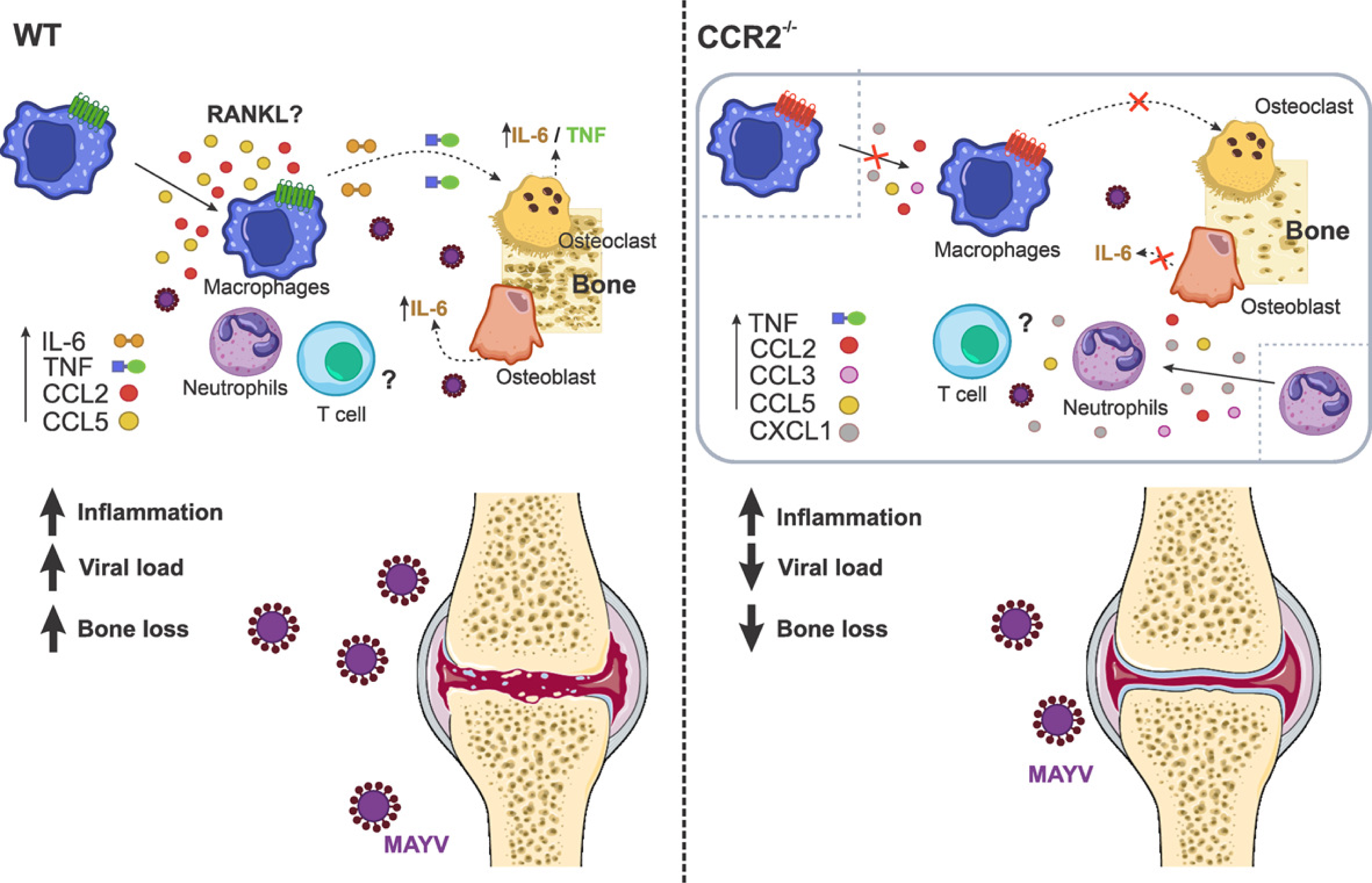
Mechanism involved in osteoclastogenesis induced by MAYV. MAYV infection induces a robust inflammatory response at the infection site in WT and CCR2^-/-^ animals, mediated by different pro-inflammatory cytokines and chemokines. This inflammatory process is responsible for tissue damage and also for bone loss. The absence of bone loss in CCR2-/-animals showed that CCR2/CCL2 signaling is essential for the migration of macrophages to the infection site. The inflammatory microenvironment induced by MAYV in different cells, promoted the differentiation of macrophages in osteoclasts by a mechanism probably independent on RANKL. It is suggested that direct action of the cytokines IL-6 and TNF are able to support the formation of new osteoclasts. The increase in osteoclasts and their activity promotes bone loss. The role of T cells in bone loss from MAYV has not been investigated in this work.

Molecular and cellular mechanisms associated to MAYV pathogenesis are poorly understood. From animal models and studies with patients, MAYV, CHIKV and RRV have been shown to induce a strong local pro-inflammatory response, characterized by the predominance of macrophages and pro-inflammatory mediators such as IL-6, TNF, INF-γ and CCL2 (8, 13, 16, 27, 28). The CCL2-CCR2 axis has been proposed to be the main route responsible for the recruitment of monocytes and macrophages during alphaviral infections. Thus, studies on the role of monocytes/macrophages and CCL2-CCR2 axis in the infection by different arthritogenic alphaviruses would allow us to understand the interaction of alphaviruses with these factors, and thus elucidate the mechanisms associated with the development of a more severe disease (7).

Our results show that infection of 4-week-old WT mice was characterized by MAYV replication and dissemination to various tissues, as well as induction of clinical disease signs, such as plantar edema and persistent articular hypernociception. These signs were accompanied by extensive inflammation of muscle tissue. These results are similar to those found in other animal models proposed for MAYV and other alphaviruses (13, 27, 29–31). The inflammatory response was characterized by local production of the chemokines CCL2 and CCL5, and the cytokines IL-6 and TNF. Interestingly, we observed high levels of CCL2 systemically, in tissues such as the maxillae, muscle and spleen. Elevated levels of the CCL5 chemokine have also been detected in the quadriceps muscle. These inflammatory mediators have been reported in other animal models for MAYV, as well as for CHIKV and RRV (8, 13, 27, 30, 32). Accordingly, patients with persistent arthralgia during MAYV infection, showed elevated levels of various inflammatory mediators, including the cytokine IL-6, which remained elevated for more than 12 months post-infection. The acute phase of the disease in these patients was marked by a high expression of CCL2, and remained elevated for months after infection (16). These mediators, which have been constantly reported in the serum of patients infected with different arthritogenic alphaviruses, indicate a common signature of the inflammatory response during infection with these viruses (6, 14, 16). Nevertheless, the role of each of these mediators in the pathogenesis of alphaviral diseases is not well established. CCL2 chemokine is important for the recruitment of monocytes/macrophages to the infection site, which are important cells in the development of myositis and arthritis (9, 10, 29). The cytokines IL-6 and TNF have been described as having an important role in tissue damage (30). In addition, CCL2 and the cytokine IL-6 have been associated with bone pathologies induced by CHIKV and RRV (11, 12, 33).

In inflammatory rheumatic diseases, inflammation is considered the main mechanism responsible for bone loss, and these cytokines have already been described as having a central role in bone loss in RA and in infection with other alphaviruses (11, 12, 34). The occurrence of bone erosion, similar to that of rheumatoid arthritis (RA), has been reported in the joint of patients infected with CHIKV. Some studies in animal models for CHIKV and RRV, as well as in osteoblast cell culture, demonstrated that the inflammation induced in the bone microenvironment triggers osteoclast-mediated bone resorption (11, 12, 33). Chen et al. (11) showed that RRV infects osteoblasts and induces IL-6 production, which in turn lead to increased expression of CCL2, to changes in the RANKL (Receptor Activator of Nuclear Factor kappa-B Ligand):OPG (Osteoprotegerin) ratio, and consequently bone loss. Blockade of IL-6 and CCL2 in animal models for RRV and CHIKV, was accompanied by a significant improvement in bone loss (9–12). In this work, we demonstrated that MAYV also has the potential to induce local bone loss. Thus, as observed for RRV and CHIKV, CCL2 is a key mediator in MAYV osteclastogenesis, given its significant increase in a systemic way in infected animals.

Arthritic disease is generally characterized by high levels of CCL2 and monocyte / macrophage infiltrates, and is also well described for alphaviral arthropathies. Given the important role of this CCL2/CCR2 pathway in the pathology of many diseases, many therapeutic agents targeting the chemokine CCL2 and the CCR2 receptor have been developed (21). Treatment of mice infected with RRV or CHIKV with bindarit, a drug that inhibits the production of CCL2, resulted in the improvement of rheumatic disease induced by these viruses (9, 10, 12). However, the absence of CCR2 has been controversial in different rheumatic diseases (20, 35). Infection of CCR2^-/-^ mice with CHIKV showed a more severe, prolonged and erosive disease that was dominated by neutrophils, with viral replication and persistence not being significantly affected (21). Interestingly, we demonstrated that CCR2^-/-^ mice infected by MAYV had a milder and delayed disease manifestation, with reduced viral loads in some tissues, fast recovery of articular hypernociception and absence of bone loss, when compared to WT. Paw edema and the peak of the inflammatory process in CCR2^-/-^ mice was delayed in relation to the WT animals. This delay can be explained by the scarcity of M2 macrophages (CCR2^+^ monocyte derivative) at the infection site, which are important cells for the resolution of inflammation and tissue regeneration. Poo et al. (21) demonstrated in their model of CCR2^-/-^ mice for CHIKV a lower upregulation of M2 macrophage markers. Interestingly, as described for CHIKV, we observed a significant increase in CXCL-1 levels, accompanied by the replacement of an infiltrate of macrophages by neutrophils in the paw of CCR2^-/-^ mice. The intense migration of CCR2^+^ monocytes to the infection sites in WT animals suggests that these cells play a central role in MAYV-mediated bone loss in these animals. Accordingly, this data is strongly confirmed by the protection against bone loss observed in knockout animals, where there is an intense reduction in the influx of monocytes. Population of resident macrophages in WT animals rapidly recovered within 7 days PI. The presence of TCD4 ^+^ and TCD8 ^+^ lymphocytes was similar in both animals.

Silencing data showed that reduction in CCL2 levels was associated with a partial improvement of clinical parameters, reinforcing the role of this pathway in the pathogenesis of MAYV. Furthermore, CCL2 silencing was also accompanied by a reduction in CCL5, CCL7 and TNF expression levels. This indicates that in the inflammatory microenvironment of MAYV infection, CCL2 secretion upregulates TNF expression, and TNF in turn stimulates the expression of CCL2 and CCL5 (36). This can result in a positive feedback loop between CCL2 and TNF. Nanki et al (37) demonstrated that TNF regulated the production of CCL2 and CCL5 in fibroblast-like synoviocytes (FLS) from patients with RA (17, 37, 38). In addition, our data also showed that CCL2 silencing modulates the expression of CCL7, an important chemokine that regulates monocytosis via CCR2 and that is often co-induced with CCL2 (36, 39). However, treatment was not enough to improve tissue damage induced by MAYV and to reduce the inflammatory infiltrate, suggesting that probably other CCR2 ligands (CCL8, CCL13 and CCL12) or other inflammatory mediators in a CCR2-independent manner can induce the migration/accumulation of cells in the target tissue (40, 41). Drevets et al. (42) demonstrated that IFN-γ is essential for influx of monocytes to the brain during systemic infection with virulent *Listeria monocytogenes* and was largely independent of a specific chemokine:receptor axis. Furthermore, the infection promoted an increase in the production of several chemokines involved in the recruitment of myeloid cells and this explains the normal influxes of Ly6C^high^ monocytes in the brain in mice deficient for CCL2, CCR1, CCR5, CXCR3 or CX3CR1 (42). This shows the importance of investigating CCR2-independent myeloid cell migration pathways in the pathogenesis of MAYV and other arthritogenic alphaviruses Interestingly, these data also show that hypernociception induced by MAYV infection is associated with the production of CCL2 and inflammatory cytokines such as TNF (43, 44). In accordance, Kwiatkowski and colleagues (44) showed that the CCL2 chemokine plays a crucial role in the development of neuropathic pain in mice (44). This reveals that CCL2 is a promising therapeutic target for the clinical treatment of MAYV-induced pain.

The long and sustained blockade of CCR2 used in this work as a strategy to prevent tissue damage and bone loss, did not present a significant protective effect. This result is similar to that found by Longobardi *et al*. in a model of OA, where they observed that long-term inhibition of 1-8 or 1-12 weeks did not prevent cartilage and bone damage, when compared with short-term treatment of 4 weeks. These findings suggest that prolonged CCR2 blockade may be ineffective in protecting against MAYV-induced joint damage and open new perspectives to explore different treatment strategies targeting CCR2 as a target molecule in MAYV-induced joint disease (45).

It has recently been shown that MAYV is capable of replicating in human osteoblasts (HOB), as well as in human chondrocytes (HC) and fibroblast-like synoviocytes (HFLS) (46). Many studies have also shown that CHIKV and RRV also replicate in cells that are important for arthritis induced by these viruses (11, 33, 46). In primary culture of WT and CCR2^-/-^ osteoclasts we observed that MAYV in MOI of 1 was not able to establish a productive infection in these cells, but induced a high production of IL-6 and TNF mainly in WT cells. TNF and IL-6 are considered the most potent osteoclastogenic cytokines and play a central role in the pathogenesis of RA (47). These cytokines can stimulate osteoclastogenesis in two ways, by a mechanism independent on RANKL or indirectly by increasing the expression of RANKL and RANK, in osteoblasts and osteoclast precursors, respectively (48, 49). Thus, according to *in vivo* and cell culture data, we suggest that the bone loss observed in WT animals may be caused by the direct action of TNF or IL-6, which is capable of inducing osteoclast formation and local osteolysis. Infection of WT and CCR2^-/-^ macrophages resulted in the production of IL-6 and TNF, respectively. Infection of BMSCs isolated from 14 d.p.d. WT and CCR2^-/-^ animals and WT osteoblasts with pharmacologically blocked CCR2 demonstrated that MAYV infection did not affect osteogenic activity of these cells, unlike what was observed by Roy et al. (50), in that CHIKV reduced ALP (Alkaline Phosphatase) activity in 9 d.p.d. infected osteoblasts. IL-6 production was induced by MAYV in WT Osteoblasts. However, in pharmacologically blocked CCR2 or CCR2^-/-^ cells, there was no production of IL-6. Taken together, these data show the importance of IL-6 production by osteoblasts in the bone microenvironment to promote deregulation of homeostasis in bone tissue, as a consequence of MAYV infection.

Overall, our results demonstrate that the absence of CCL2/CCR2 signaling minimized the disease induced by MAYV and mitigated MAYV-driven bone loss. In this way, we were able to demonstrate one of the mechanisms underlying MAYV-induced bone pathology and consequently the role played by CCL2/CCR2 axis as a potent therapeutic target.

## Material and Methods

### Cells, virus and titration

Mayaro virus strain is a human isolate from Peru (in 2001) obtained from the World Reference Center for Emerging Viruses and Arboviruses at the University of Texas Medical Branch. MAYV stocks were produced on Vero cells CCL-81 from BCRJ (African green monkey kidney cell line). Briefly, Vero cells were grown in Dulbecco’s Modifed Eagle’s Medium-DMEM medium (Cultilab, Brazil) supplemented with 10% inactivated fetal bovine serum (FBS; Cultilab, Brazil) and 1% Penicillin/Steptamycin/Glutamine (GIBCO), and kept in a humidified incubator at 37°C with 5% CO_2_ atmosphere for 4 days. Cell supernatant was collected, centrifuged (2000rpm for 10 min) and then concentrated using a Vivacell 100 centrifugal concentrator (Sartorius). MAYV titration was performed as described in Mota & Costa 2020. Briefly, Vero cells were plated in 24-well plates and infected with 10-fold serial dilutions. Plates were incubated for 72 hours (37°C with 5% CO_2_ atmosphere), and then fixed with 10% formaldehyde and stained with 1% violet crystal. Plaques were counted by eye.

### Ethical Statement

This study was carried out in strict accordance with the Brazilian Government’s ethical and animal experiments regulations (Law 11794/2008). The experimental protocol was approved by the Committee on the Ethics of Animal Experiments of the Universidade Federal de Minas Gerais (CEUA/UFMG, Protocol Number 160/2018). All surgeries were performed under ketamine/xylazine anesthesia, and all efforts were made to minimize animal suffering.

### Infection of animals

Four-week-old male C57BL/6J mice (wild-type-WT) were obtained from the Biotério Central of the Universidade Federal de Minas Gerais - UFMG. CCR2 receptor-deficient male mice (CCR2^-/-^) were obtained from the Biotério de Imunofarmacologia of UFMG. WT and CCR2^-/-^ mice were infected on the right footpad with 1×10^6^ PFU of MAYV, with a final volume of 30μl. Mock-infected mice received 30μl of PBS 1X.

### Evaluation of disease parameters

Mice were observed up to 21 days after infection. During this period, every 24 hours, mice were evaluated for signs of disease, such as weight loss, piloerection and paw edema. Paw edema was quantified with the aid of a caliper and measurements were performed on the infected paw, at times points 0, 1, 3, 5, 7, 10, 14 and 21 days after infection.

### Hypernociception assessment by a modified electronic pressure-meter test von Frey method

Hypernociception was assessed as described by Costa et al., 2012. Briefly, mice were placed in acrylic boxes with a non-malleable wire floor for 15 minutes for acclimatization. Then, with the aid of a pressure transducer adapted to a standard large (0.5 mm^2^) polypropylene tip (INSIGTH Instruments, Ribeirao Preto, SP, Brazil), an increasing pressure was exerted to the right hindpaw of MOCK or MAYV infected mice until induce the flexion of the knee joint, followed by paw withdraw. Upon the flexion-elicited withdrawal threshold, the intensity of the pressure was automatically recorded. Two measurements were performed per each animal at each time analyzed and the median was used. Baseline measurements of all groups were initially performed before infection, and the remaining measurements were taken 1, 3, 7, 14, 21 and 28 days post-infection.

### ELISA assay

To assess the profile of cytokines and chemokines during infection, plasma and tissues of interest were collected at different post-infection times and assessed by ELISA using commercially available antibodies and according to the procedures supplied by the manufacturer (R&D Systems, Minneapolis). All samples were stored at -20°C until processing. Tissues were macerated in cytokine buffer, centrifuged at 10,000 rpm for 10min at 4°C. The supernatant was collected and stored until the ELISA was performed. Results are expressed as pg/mL or pg/100 mg of tissue. The detection limit of the ELISA assays was in the range of 4–8 pg/ml.

### Micro-CT analysis

After euthanasia, mice right hind paw was removed, fixed in 10% formaldehyde and then transferred to 70% alcohol, where they were kept until analysis in the micro-CT. Samples were scanned on a compact desktop micro-CT scanner (SkyScan 1174, Bruker micro-CT, Belgium) operating at 50 kV of source voltage, 800 µA source current, 14.59 µm pixel size and a 0.5mm Al filter. Samples were attached to a stage that rotated 180° with images acquired every 0.7°. The acquired shadow projections (16-bit TIFF format) were further reconstructed into 2D slices using the NRecon software interface. 3D analyses were performed on SkyScan CTAn tool, and 3D models were observed SkyScan CTVol interface.

### Histopathological analysis

After euthanasia, tissues of interest were removed and fixed in 10% formaldehyde. For analysis of ankle joint, samples were decalcified in 10% EDTA, pH 7.0. Then tissues were embedded in paraffin, and sections of 5μm were performed. Analysis of tissue damage and inflammatory infiltrate were performed on mice footpad samples stained hematoxylin and eosin (H & E) slides. The samples were evaluated by a blinded pathologist using a score on inflammatory infiltrate and damage to muscle architecture, in a scale varying from no lesion to mild, moderate, intense and very intense lesion. The data are presented as percentage of mice presenting each lesion degree in each experimental group with a contingency table. To assess osteoclastogenesis, selected slides from the ankle joint with 21 days PI were stained for tartrate-resistant acid phosphatase (TRAP; Sigma-Aldrich, Saint Louis, MO, USA) and counterstained with hematoxylin according to the manufacturer’s instructions.

### Footpad cell isolation

Footpad and ankle joint from control and MAYV infected mice, were collected at 1 and 7 days post-infection. For isolation of cells from the plantar pad and ankle, tissues were removed and placed in 1 mL of digestion medium containing Collagenase VIII (1 mg / mL; Sigma-Aldrich) in complete DMEM medium. Tissues were kept for 1h in the water bath at 37°C. Then, the digested tissues were deposited in 40μm cell strainer and were added 3 ml complete DMEM medium. Digested tissues were macerated against the cell strainer with a 3mL syringe plunger, making circular movements to release the largest number of cells in the medium. Cells were then centrifuged at 1,200 rpm for 10 minutes at 4°C, and resuspended in 1mL of complete medium. Cell viability was assessed with Trypan Blue.

### Phenotyping of leukocytes

Cells from lymph node and paw were plated on a 96-well plate with a U-shaped bottom to perform staining. First, the plate was centrifuged at 2,000 rpm for 10 min at 4°C to remove medium. Cells were resuspended in 20μl of Fc-blocker (BD Bioscienses) for 15 minutes. Staining was performed using fluorescently conjugated antibodies anti-CD3, - CD4, -CD8, -CD69, -CD44, -CD62L, -CCR2, -CD11c, -F4/80, -CD11b, -Ly6G, -Ly6C, - MHCII, -CD80, for 30 min. Then, 200 μl of 1% BSA was added and the cells were centrifuged to remove antibody excess. Cells were resuspended in 200 μl of 1% BSA and the data was acquired using a BD FACS CANTO II and analysis performed using the FlowJo software.

### Reverse transcription and Real Time PCR (RT-qPCR) analysis

WT mice were infected and after euthanasia paw were removed and stored at -80°C until processing. For total RNA extraction, samples were first mashed in liquid nitrogen and followed by Trizol extraction according to the manufacturer’s instructions. The reverse transcription was performed with random primers and following the guidelines of iScript cDNA Synthesis Kit-Bio-Rad. The real time PCR reaction was performed as follows: 2 µl of RT product (50ng concentration), 1 µl of each primer, 5 µl of Syber Green (qPCR Master Mix - Bio-Rad) and 2 µl of water. The conditions for PCR were as follows: 95 C – 2 min, followed by 40 cycles of 95 C – 15 s; 60 C – 1 min, and 72 C – 20 s. The efficiency/slope obtained values of all investigated genes were close to the optimal values required for the 2(-Delta Ct) analysis (51). Ct values were recorded for each gene and the results regarding the genes of interest were normalized to the results obtained with the internal control genes, *RPS18 and PPIA*. The ΔΔCT values were calculated and the results were expressed as fold increase, as described by Livak and Schmittgen (52). Sequence of primers:

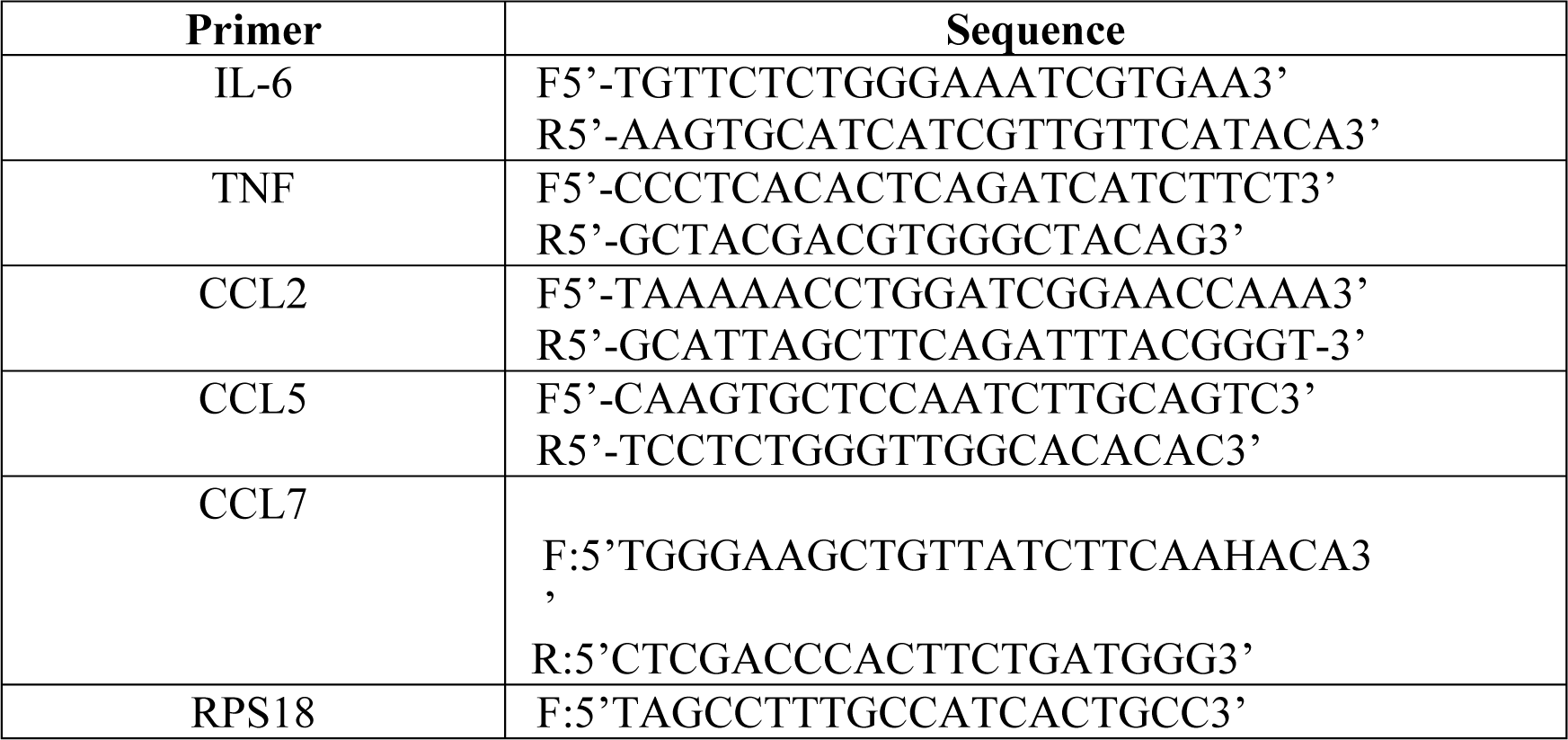
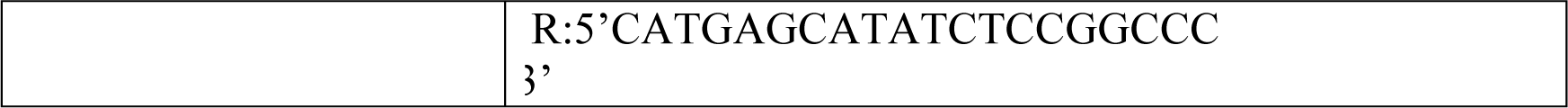

### *In vivo* silencing of the CCL2 chemokine with siRNA

Nanoparticles carrying siRNA-MCP1 (NP-siRNA-CCL2) were synthesized according to Krohn-Grimberghe et al (2020). For the silencing of the CCL2 chemokine, 4-week-old animals of the C57BL/6J strain were divided into 3 experimental groups: i) sham: received only PBS; ii) MAYV: infected only and iii) MAYV+SiRNA: infected and treated. The animals in the treated group received 1mg/kg intravenously. The treatment strategy included a pre-treatment for 3 days, every 24h, before infection and post-infection treatments on days 2 and 5. After infection, the animals were evaluated for disease parameters, such as edema and hypernociception at defined time points. On day 7 PI, the animals were sacrificed and the tissues of interest were removed for the following analyses, viral load, histology, ELISA, flow cytometry and gene expression.

### *In vivo* CCR2 antagonism

For systemic CCR2 receptor blockade, mice were treated with RS504393 (Sigma, Aldrich), a CCR2 antagonist. Treatment started 24 h before infection with MAYV. Every 24 h until the end of the experiment, the mice received intraperitoneal injections of 4 mg/kg of RS504393 (dissolved in DMSO at 10 mM and diluted in PBS for application, groups RS 504393 and MAYV+RS504393). Untreated groups (NI group) received 100 μl of sterile PBS or DMSO 10 mM in PBS (DMSO group).

### Cell culture

For primary osteoblast, osteoclast and macrophages cultures, bone marrow cells derived from WT and CCR2^-/-^ mice were isolated and kept in culture for growth, in α-MEM medium with 10% FBS at 37°C and 5% CO2. For osteoblast culture the cells were then plated into 96 or 24-well plates, and differentiation was induced with osteogenic medium (α-MEM medium 10% plus 100µM ascorbic acid and 1.8nM potassium phosphate – KH_2_PO_4_) for 14 days. To evaluate the effect of blocking CCR2 in osteoblast culture, cells were treated 1h before infection with the antagonist RS504393 (300nM per well). Every two days the medium was replaced with new medium. After differentiation, cells were infected with MAYV and cell supernatant collected at different times points post-infection. Several analyzes were performed, including viral titration, ELISA, cell viability (MTT) and alkaline phosphatase labeling. For macrophage culture the cells were then plated into a 96-well plate, and differentiation was induced with 50ng/ml of M-CSF for 3 days. After differentiation, cells were infected with MAYV and cell supernatant collected at different times points post-infection for analysis. For osteoclast culture the cells were then plated into a 96-well plate, and differentiation was induced with 50ng/ml of M-CSF and 50ng/ml of RANKL for 5 days.

### Statistical analysis

All analyzes were performed using the GraphPad PRISM software 6.0 (GraphPad Software, USA). Results with a p <0.05 were considered significant. Differences were compared using one or two-way analysis of variance (ANOVA) followed by Tukey’s multiple comparisons test or Unpaired t test.

## Acknowledgments

The authors would like to thank Ilma Marçal, Tania Colina, Frankcineia Assis, and Gilvania Santos for the technical support. This work received financial support from the National Institute of Science and Technology in Dengue and Host-microorganism Interaction (INCT dengue), a program funded by The Brazilian National Science Council (CNPq, Brazil) and Minas Gerais Foundation for Science (FAPEMIG, Brazil) and the Coordenação de Aperfeiçoamento de Pessoal de Nível Superior (CAPES, Brazil).

## Author Contributions

Franciele Martins Santos: Conceptualization, Data curation, Formal analysis, Investigation, Methodology, Visualization, Writing – original draft

Victor Rodrigues de Costa Melo: Investigation and Methodology

Simone de Araújo: Data curation, Investigation and Methodology

Carla Daiane Ferreira de Sousa: Data curation, Investigation and Methodology

Thaiane Pinto Moreira: Data curation, Investigation and Methodology

Matheus Rodrigues Gonçalves: Investigation and Methodology

Breno Rocha Barrioni: Methodology

Anna Clara Paiva Menezes dos Santos: Methodology

Heloísa A Seabra: Data curation, Investigation, Methodology, review and editing

Pedro A C Costa: Data curation, Investigation and Methodology, review and editing.

Breno Rocha Barrioni: Investigation and Methodology

Paula Bargi-Souza: RT-qPCR standardization, Data curation, analysis of the results, Writing - review and editing

Renato Santana Aguiar: Resources

Marivalda de Magalhães Pereira: Methodology and Resources Maurício Lacerda Nogueira: Resources

Danielle Souza da Glória: Resources

Pedro P G Guimarães: Investigation, Resources and Writing – review and editing

Mauro Martins Teixeira: Funding acquisition and Resources

Celso Martins Queiroz-Junior: Data curation, Investigation, Supervision, Visualization, Writing - review and editing

Vivian Vasconcelos Costa: Funding acquisition, Investigation, Project administration, Supervision, Validation, Visualization, Writing – review and editing

## References

1. Y. Acosta-Ampudia, et al., Mayaro: an emerging viral threat? Emerging Microbes & Infections 7, 1–11 (2018).

2. M. T. de O. Mota, M. R. Ribeiro, D. Vedovello, M. L. Nogueira, Mayaro virus: a neglected arbovirus of the Americas. Future Virology 10, 1109–1122 (2015).

3. E. S. Halsey, et al., Mayaro Virus Infection, Amazon Basin Region, Peru, 2010–2013. Emerg. Infect. Dis. 19 (2013).

4. C. Theilacker, et al., Prolonged polyarthralgia in a German traveller with Mayaro virus infection without inflammatory correlates. BMC Infect Dis 13, 369 (2013).

5. C. A. D. Slegers, et al., Persisting arthralgia due to Mayaro virus infection in a traveler from Brazil: Is there a risk for attendants to the 2014 FIFA World Cup? Journal of Clinical Virology 60, 317–319 (2014).

6. D. Tappe, et al., Sustained Elevated Cytokine Levels during Recovery Phase of Mayaro Virus Infection. Emerg. Infect. Dis. 22, 750–752 (2016).

7. H. Mostafavi, E. Abeyratne, A. Zaid, A. Taylor, Arthritogenic Alphavirus-Induced Immunopathology and Targeting Host Inflammation as A Therapeutic Strategy for Alphaviral Disease. Viruses 11, 290 (2019).

8. B. A. Lidbury, et al., Macrophage-Derived Proinflammatory Factors Contribute to the Development of Arthritis and Myositis after Infection with an Arthrogenic Alphavirus. J INFECT DIS 197, 1585–1593 (2008).

9. N. E. Rulli, et al., Amelioration of alphavirus-induced arthritis and myositis in a mouse model by treatment with bindarit, an inhibitor of monocyte chemotactic proteins. Arthritis Rheum 60, 2513–2523 (2009).

10. N. E. Rulli, et al., Protection From Arthritis and Myositis in a Mouse Model of Acute Chikungunya Virus Disease by Bindarit, an Inhibitor of Monocyte Chemotactic Protein-1 Synthesis. The Journal of Infectious Diseases 204, 1026–1030 (2011).

11. W. Chen, S.-S. Foo, R. W. Li, P. N. Smith, S. Mahalingam, Osteoblasts from osteoarthritis patients show enhanced susceptibility to Ross River virus infection associated with delayed type I interferon responses. Virol J 11, 189 (2014).

12. W. Chen, et al., Arthritogenic alphaviruses: new insights into arthritis and bone pathology. Trends in Microbiology 23, 35–43 (2015).

13. F. M. Santos, et al., Animal model of arthritis and myositis induced by the Mayaro virus. PLoS Negl Trop Dis 13, e0007375 (2019).

14. T.-S. Teng, et al., A Systematic Meta-analysis of Immune Signatures in Patients With Acute Chikungunya Virus Infection. J Infect Dis. 211, 1925–1935 (2015).

15. K. C. Haist, K. S. Burrack, B. J. Davenport, T. E. Morrison, Inflammatory monocytes mediate control of acute alphavirus infection in mice. PLoS Pathog 13, e1006748 (2017).

16. F. W. Santiago, et al., Long-Term Arthralgia after Mayaro Virus Infection Correlates with Sustained Pro-inflammatory Cytokine Response. PLoS Negl Trop Dis 9, e0004104 (2015).

17. M. Gschwandtner, R. Derler, K. S. Midwood, More Than Just Attractive: How CCL2 Influences Myeloid Cell Behavior Beyond Chemotaxis. Front. Immunol. 10, 2759 (2019).

18. R. E. Miller, et al., CCR2 chemokine receptor signaling mediates pain in experimental osteoarthritis. Proc. Natl. Acad. Sci. U.S.A. 109, 20602–20607 (2012).

19. C.-Y. Chen, et al., Enhancement of CCL2 expression and monocyte migration by CCN1 in osteoblasts through inhibiting miR-518a-5p: implication of rheumatoid arthritis therapy. Sci Rep 7, 421 (2017).

20. H. Raghu, et al., CCL2/CCR2, but not CCL5/CCR5, mediates monocyte recruitment, inflammation and cartilage destruction in osteoarthritis. Ann Rheum Dis 76, 914–922 (2017).

21. Y. S. Poo, et al., CCR2 Deficiency Promotes Exacerbated Chronic Erosive Neutrophil-Dominated Chikungunya Virus Arthritis. J Virol 88, 6862–6872 (2014).

22. P. J. Hotez, K. O. Murray, Dengue, West Nile virus, chikungunya, Zika—and now Mayaro? PLoS Negl Trop Dis 11, e0005462 (2017).

23. I. Assunção-Miranda, C. Cruz-Oliveira, A. T. Da Poian, Molecular Mechanisms Involved in the Pathogenesis of Alphavirus-Induced Arthritis. BioMed Research International 2013, 1– 11 (2013).

24. D. Damjanovic, C.-L. Small, M. Jeyananthan, S. McCormick, Z. Xing, Immunopathology in influenza virus infection: Uncoupling the friend from foe. Clinical Immunology 144, 57– 69 (2012).

25. A. Culshaw, J. Mongkolsapaya, G. R. Screaton, The immunopathology of dengue and Zika virus infections. Curr Opin Immunol 48, 1–6 (2017).

26. J. N. Gustine, D. Jones, Immunopathology of Hyperinflammation in COVID-19. The American Journal of Pathology 191, 4–17 (2021).

27. J. Gardner, et al., Chikungunya Virus Arthritis in Adult Wild-Type Mice. J Virol 84, 8021– 8032 (2010).

28. L. F. P. Ng, Immunopathology of Chikungunya Virus Infection: Lessons Learned from Patients and Animal Models. Annu. Rev. Virol. 4, 413–427 (2017).

29. T. E. Morrison, et al., Characterization of Ross River Virus Tropism and Virus-Induced Inflammation in a Mouse Model of Viral Arthritis and Myositis. J Virol 80, 737–749 (2006).

30. C. M. Figueiredo, et al., Mayaro Virus Replication Restriction and Induction of Muscular Inflammation in Mice Are Dependent on Age, Type-I Interferon Response, and Adaptive Immunity. Front. Microbiol. 10, 2246 (2019).

31. M. T. de O. Mota, et al., In-depth characterization of a novel live-attenuated Mayaro virus vaccine candidate using an immunocompetent mouse model of Mayaro disease. Sci Rep 10, 5306 (2020).

32. L. J. Herrero, et al., Critical role for macrophage migration inhibitory factor (MIF) in Ross River virus-induced arthritis and myositis. Proc. Natl. Acad. Sci. U.S.A. 108, 12048–12053 (2011).

33. M. Noret, et al., Interleukin 6, RANKL, and Osteoprotegerin Expression by Chikungunya Virus-Infected Human Osteoblasts. Journal of Infectious Diseases 206, 455–457 (2012).

34. F. Coury, O. Peyruchaud, I. Machuca-Gayet, Osteoimmunology of Bone Loss in Inflammatory Rheumatic Diseases. Front. Immunol. 10, 679 (2019).

35. M. P. Quinones, et al., The complex role of the chemokine receptor CCR2 in collagen-induced arthritis: implications for therapeutic targeting of CCR2 in rheumatoid arthritis. J Mol Med 83, 672–681 (2005).

36. P. Proost, A. Wuyts, J. van Damme, Human monocyte chemotactic proteins-2 and -3: structural and functional comparison with MCP-1. J Leukoc Biol 59, 67–74 (1996).

37. T. Nanki, K. Nagasaka, K. Hayashida, Y. Saita, N. Miyasaka, Chemokines Regulate IL-6 and IL-8 Production by Fibroblast-Like Synoviocytes from Patients with Rheumatoid Arthritis. J Immunol 167, 5381–5385 (2001).

38. E. Neumark, O. Sagi-Assif, B. Shalmon, A. Ben-Baruch, I. P. Witz, Progression of mouse mammary tumors: MCP-1-TNF? cross-regulatory pathway and clonal expression of promalignancy and antimalignancy factors. Int. J. Cancer 106, 879–886 (2003).

39. S. V. Bardina, et al., Differential Roles of Chemokines CCL2 and CCL7 in Monocytosis and Leukocyte Migration during West Nile Virus Infection. J.I. 195, 4306–4318 (2015).

40. J. J. Haringman, P. P. Tak, Chemokine blockade: a new era in the treatment of rheumatoid arthritis? Arthritis Res Ther 6, 93 (2004).

41 C. Shi, E. G. Pamer, Monocyte recruitment during infection and inflammation. Nat Rev Immunol 11, 762–774 (2011).

42 D. A. Drevets, M. J. Dillon, J. E. Schawang, J. A. Stoner, P. J. M. Leenen, IFN-γ triggers CCR2-independent monocyte entry into the brain during systemic infection by virulent Listeria monocytogenes. Brain, Behavior, and Immunity 24, 919–929 (2010).

43. A. Hess, et al., Blockade of TNF-α rapidly inhibits pain responses in the central nervous system. Proc. Natl. Acad. Sci. U.S.A. 108, 3731–3736 (2011).

44 K. Kwiatkowski, et al., Bidirectional Action of Cenicriviroc, a CCR2/CCR5 Antagonist, Results in Alleviation of Pain-Related Behaviors and Potentiation of Opioid Analgesia in Rats With Peripheral Neuropathy. Front. Immunol. 11, 615327 (2020).

45 L. Longobardi, et al. Role of the C-C chemokine receptor-2 in a murine model of injury-induced osteoarthritis. Osteoarthritis Cartilage. 25(6):914–925, 2017.

46. M. Bengue, et al., Mayaro Virus Infects Human Chondrocytes and Induces the Expression of Arthritis-Related Genes Associated with Joint Degradation. Viruses 11, 797 (2019).

47. W. Phuklia, et al., Osteoclastogenesis induced by CHIKV-infected fibroblast-like synoviocytes: A possible interplay between synoviocytes and monocytes/macrophages in CHIKV-induced arthralgia/arthritis. Virus Research 177, 179–188 (2013).

48. M. Ponzetti, N. Rucci, Updates on Osteoimmunology: What’s New on the Cross-Talk Between Bone and Immune System. Front. Endocrinol. 10, 236 (2019).

49. O. Kudo, et al., Interleukin-6 and interleukin-11 support human osteoclast formation by a RANKL-independent mechanism. Bone 32, 1–7 (2003).

50 E. Roy, W. Shi, B. Duan, S. P. Reid, Chikungunya Virus Infection Impairs the Function of Osteogenic Cells. 5, 12 (2020).

51. A.-A. Dussault, M. Pouliot, Rapid and simple comparison of messenger rna levels using real-time PCR. Biol. Proced. Online 8, 1–10 (2006).

52. K. J. Livak, T. D. Schmittgen, Analysis of Relative Gene Expression Data Using Real-Time Quantitative PCR and the 2−ΔΔCT Method. Methods 25, 402–408 (2001).

